# Hierarchical disruption in the cortex of anesthetized monkeys as a new signature of consciousness loss

**DOI:** 10.1101/2020.06.04.133538

**Authors:** Camilo Miguel Signorelli, Lynn Uhrig, Morten Kringelbach, Bechir Jarraya, Gustavo Deco

**Affiliations:** Department of Computer Science, University of Oxford, UK; Cognitive Neuroimaging Unit, Institut National de la Santé et de la Recherche Médicale U992, France; Center for Brain and Cognition, Computational Neuroscience Group, Universitat Pompeu Fabra, Spain; Commissariat à l’Énergie Atomique et aux Energies Alternatives, Direction de la Recherche Fondamentale, NeuroSpin Center, France; Department of Anesthesiology and Critical Care, Necker Hospital, University Paris Descartes, France; Department of Anesthesiology and Critical Care, Sainte-Anne Hospital, University Paris Descartes, France; Center for Music in the Brain, Department of Clinical Medicine, Aarhus University & The Royal Academy of Music Aarhus/Aalborg, Denmark; Centre for Eudaimonia and Human Flourishing, University of Oxford, UK; Department of Psychiatry, University of Oxford, UK; Neurosurgery Department, Foch Hospital, Suresnes, France; University of Versailles Saint-Quentin-en-Yvelines, Université Paris-Saclay, France; Department of Information and Communication Technologies, Universitat Pompeu Fabra, Spain; Institució Catalana de la Recerca i Estudis Avançats, Spain; Department of Neuropsychology, Max Planck Institute for Human Cognitive and Brain Sciences, Germany; Turner Institute for Brain and Mental Health, Monash University, Melbourne, Australia

**Keywords:** Anesthesia, Integration, Ignition, Measures of Consciousness, IIT, GNW

## Abstract

Anesthesia induces a reconfiguration of the repertoire of functional brain states leading to a high function-structure similarity. However, it is unclear how these functional changes lead to loss of consciousness. Here we suggest that the mechanism of conscious access is related to a general dynamical rearrangement of the intrinsic hierarchical organization of the cortex. To measure cortical hierarchy, we applied the Intrinsic Ignition analysis to resting-state fMRI data acquired in awake and anesthetized macaques. Our results reveal the existence of spatial and temporal hierarchical differences of neural activity within the macaque cortex, with a strong modulation by the depth of anesthesia and the employed anesthetic agent. Higher values of Intrinsic Ignition correspond to rich and flexible brain dynamics whereas lower values correspond to poor and rigid, structurally driven brain dynamics. Moreover, spatial and temporal hierarchical dimensions are disrupted in a different manner, involving different hierarchical brain networks. All together suggest that disruption of brain hierarchy is a new signature of consciousness loss.

## 1. Introduction

Recent studies suggest dynamical disruptions on brain activity during general anesthesia, sleep and disorders of consciousness (Dehaene & Changeux, 2011; Mashour et al., 2020). Nevertheless, whatever the level of consciousness, the resting-state brain activity displays highly organized coherent networks (Biswal, Yetkin, Haughton, & Hyde, 1995; Buckner, Andrews-Hanna, & Schacter, 2008; Fox et al., 2005; Fransson, 2006; Vincent et al., 2007). Examples are anticorrelated networks still present under anesthesia (Boveroux et al., 2010; Vincent et al., 2007) and early stages of sleep (Fukunaga et al., 2006; Picchioni et al., 2008). Evidence suggests that anesthesia modulates the strength of functional connectivity (Martuzzi, Ramani, Qiu, Rajeevan, & Constable, 2010; Barttfeld et al., 2015; Boveroux et al., 2010; Schrouff et al., 2011). In Barttfeld et al. (Barttfeld et al., 2015), dynamical resting-state analyses of fMRI data acquired in awake and propofol anesthetized macaques, indicate that during the awake state, the brain activity at rest displays a rich repertoire of flexible functional patterns that is independent of the underlying anatomical connectivity. Conversely, during anesthesia-induced loss of consciousness, the resting-state brain activity is shifted toward a poor repertoire of rigid functional patterns with higher similarity to structural connectivity. A finding that was generalized to different anesthetic agents (Uhrig et al., 2018) and also applied to classify different categories of chronic loss of consciousness (Demertzi et al., 2019). This dynamical disruption at long-distance networks might be the common fingerprint of all different types of loss of consciousness (anesthesia-induced, injuries-induced loss of consciousness and sleep).

Unfortunately, it is still unclear how these functional disruptions are induced and if they are causal or consequence of other factors. We hypothesised these dynamical disruptions are due to the breakdown of the hierarchical organization of the cortex and cortico-sub-cortical networks (Mesulam, 1998). Independently of the molecular pathways of different anesthetics, stages of sleep or localization/types of brain injuries, if this is enough to disturb the hierarchical structure of the conscious brain, it will lead to a loss of consciousness. Differently than previous hierarchical auditory regularities studied in awake and anesthetized macaques (Bekinschtein et al., 2009; Uhrig, Janssen, Dehaene, & Jarraya, 2016), the causal driven forces that generate similar global disruptions would correspond to any local or global disturbance with enough power to reorganize the network hierarchy. These disruptions become a common mechanism for loss of consciousness, at the same time as saving the specificity of different impairments (Sherrington, 1906).

To investigate these brain mechanisms of consciousness loss, a newly introduced measure called Intrinsic Ignition, together with general anesthesia, offer a unique opportunity to quantify the hierarchy of neural activity and its disruptions (Deco & Kringelbach, 2017). On the one hand, this is possible through the massive modulation of both arousal and conscious access (i.e. awareness) by elective pharmacological drugs, called anesthetic agents. Different anesthetics, with different pharmacological and molecular pathways, generate comparable dynamical disruption (Uhrig et al., 2018). On the other hand, Intrinsic Ignition quantifies the neural propagation activity in space and time, from one region to other areas of the brain (Deco et al., 2017).

Intrinsic Ignition combines simplified versions of integration from the “integrated information theory (IIT)” (Deco, Tononi, Boly, & Kringelbach, 2015; Tononi & Koch, 2015) and broadcasting from “the global neuronal workspace theory (GNW)” (Dehaene & Changeux, 2011). Moreover, Intrinsic Ignition is complementary to the concept of ignition/broadcasting from the GNW theory. The former being considered intrinsic (due to internal interactions) and the later extrinsic (due to external stimuli) (Van Vugt et al., 2018). Broadcasting, as the first stage of neural processing and integration as the second stage, are combined into one measure of brain activity (Deco & Kringelbach, 2017). Intrinsic Ignition uses the graph theory to define integration. Integration is the accumulative and averaged value of the maximal path in a network at different states, computing the value among spatial areas and time evolution. This measure quantifies different modes of consciousness and estimates the type of hierarchical organization for different conditions. At spontaneous waking brain activity, analyses using Intrinsic Ignition suggest that the brain organization is maximally hierarchical, but not uniformly graded (Deco & Kringelbach, 2017; Deco, Tagliazucchi, Laufs, Sanjuán, & Kringelbach, 2017). Intrinsic Ignition has been applied to compare awake versus sleep conditions (Deco & Kringelbach, 2017; Deco et al., 2017) and normal subjects versus meditators (Escrichs et al., 2019).

Here, we measured Intrinsic Ignition of cortical areas from awake and anesthetized macaques, with unique access to six experimental conditions (awake, ketamine, light/deep propofol, light/deep sevoflurane anesthesia) (Barttfeld et al., 2015; Uhrig et al., 2018). Intrinsic Ignition of cortical areas assessed with fMRI reveals spatial-temporal hierarchical differences and allows clustering anaesthetics. Finally, Intrinsic Ignition, as a unifying concept across theoretical frameworks of consciousness, quantifies the dynamical disruption in terms of hierarchical organization arrangement and defines a multidimensional signature of consciousness.

## 2. Results

### 2.1 Dynamical differences among Anesthetics

The analyzed data here corresponds to 119 runs. An example of the time series for one subject in the awake condition is plotted in Figure 1a. The probability density (density distribution) of fMRI values is plotted after a normalization procedure (using z-score). These plots suggest the use of non-parametric statistical tests since the center of the data distributions present close to zero mean, but seemingly different variance. The Kolmogorov–Smirnov test finds statistical differences among all conditions (p<0.001, CI reported in Captions Figure 1b). The FC matrix plotted as the average among subjects also supports dynamical differences among conditions (Figure 1c).

**Figure 1.**
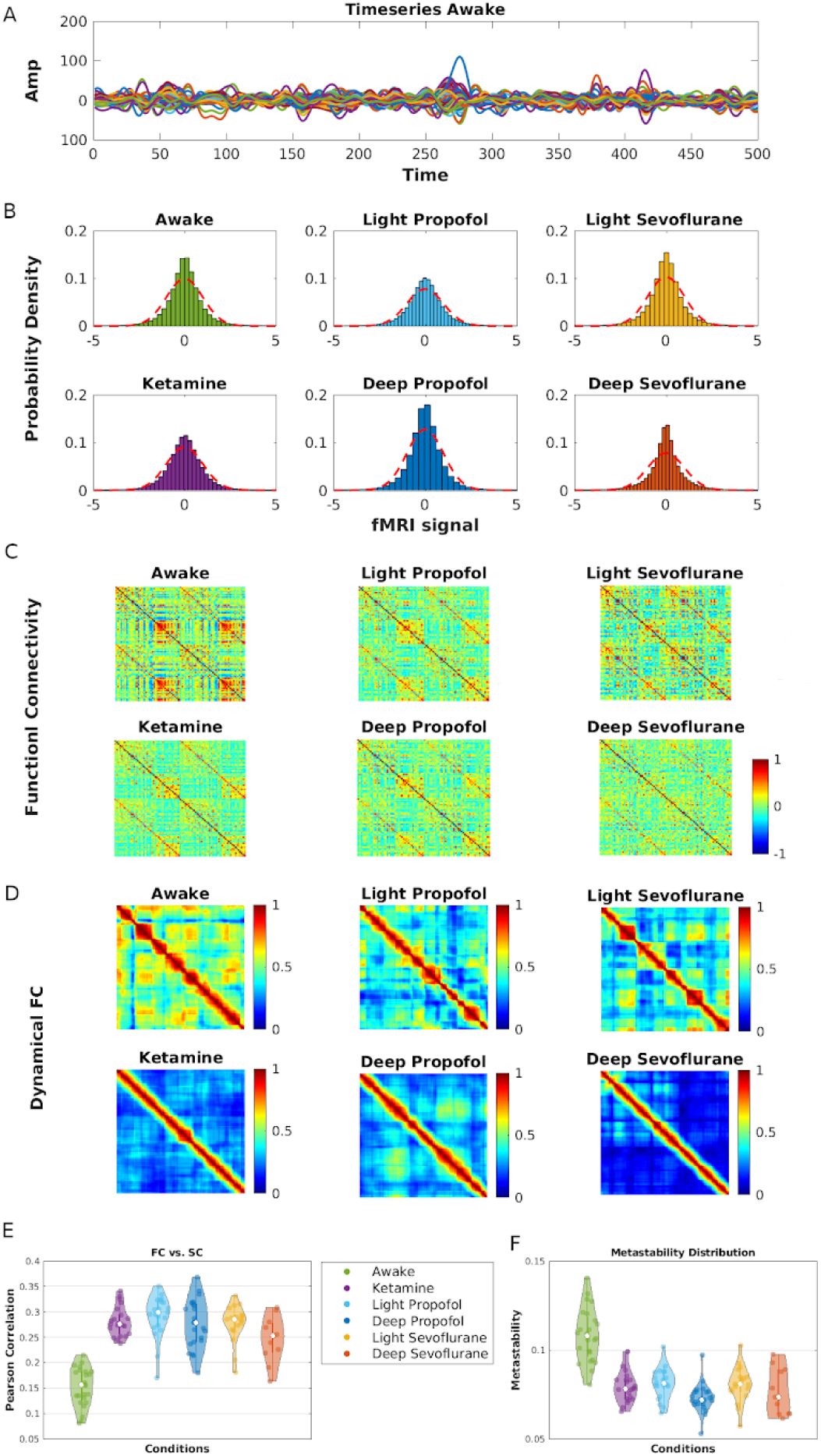
Dynamical Analysis. **A)** Example of time series for one monkey, awake condition, 500-time points with a 2400 ms repetition time (TR). **B)** Distribution plots of the fMRI signal for each condition. All conditions are significantly different (Kolmogorov–Smirnov test p<0.001, awake CI [0.0062 −0.0332], ketamine [0.0027 −0.0021], light propofol CI [0.0025 −0.0026], deep propofol CI [0.0033 −0.0029], light sevoflurane CI [0.0032 −0.0058], deep sevoflurane anesthesia CI [0.0061 −0.0057]). **C)** Functional connectivity matrices (FC) for each condition, CoCoMac, 82 cortical regions. **D)** Example of dynamical functional connectivity (dFC) for one subject in each condition. **E)** Pearson Correlation between FC and Structural connectivity (SC) for each subject and plotted as a violin plot, see methods, and (Hintze & Nelson, 1998). Awake condition is significantly different than the other conditions (Kolmogorov–Smirnov test p<0.001, awake CI [0.1671 0.1369], ketamine CI [0.2926 0.2687], light propofol CI [0.3093 0.2738], deep propofol CI [0.2957 0.2568], light sevoflurane CI [0.2943 0.2601], deep sevoflurane CI [0.2725 0.2196]), while other conditions are not statistically different (p>0.01). **F)** Metastability is higher in the awake state than in the anesthesia conditions (Kolmogorov–Smirnov test p<0.001, awake CI [0.1143 0.1024], ketamine CI [0.0835 0.0759], light propofol CI [0.0849 0.0775], deep propofol CI [0.0762 0.0697], light sevoflurane CI [0.0853 0.0759], deep sevoflurane CI [0.0844 0.0687]).

These differences also appear when plotting the dynamical functional connectivity (dFC) (Figure 1d). Awake (followed by light propofol and light sevoflurane sedation) seems to present more correlated activity among functional matrices across time than the deep anesthesia conditions. Deep propofol anesthesia is slightly more activated than ketamine and deep sevoflurane. To quantify these qualitative differences a Pearson correlation between the FC per subject and the structural connectivity (SC, CoCoMac) is performed (Figure 1e). It reveals that the awake state has a lower correlation value, indicating that functional activity is farther from SC, than other conditions (Kolmogorov–Smirnov test, p<0.001). The violin plots in Figure 1e also demonstrate differences in the type of distributions. Light propofol sedation has the highest mean correlation value, however, anesthetics are not differentiated in terms of statistical tests (p>0.01). Quantifying the dynamical variability through metastability, as the standard deviation of the Kuramoto’s order parameter (synchrony), shows the awake condition with higher values of metastability (Figure 1f, Kolmogorov–Smirnov test, p<0.001), light propofol and light sevoflurane sedation slightly higher than ketamine and deep sedations, but not statistically significant. Deep propofol anesthesia has a slightly lower value than ketamine anesthesia (Kolmogorov–Smirnov test, p=0.01), as well as light propofol (p<0.001) and light sevoflurane anesthesia (p=0.0011). Other conditions do not present major statistical differences in terms of metastability (p>0.01).

These dynamical analyses support and replicate previous results suggesting disruption of the dynamical functional organization under anesthesia (Barttfeld et al., 2015; Uhrig et al., 2018). However, they do not distinguish among anesthetics or quantify the degree of disruption in terms of hierarchical organization. Therefore, extra analyses may offer a complementary picture.

### 2.2 Intrinsic Ignition and global Hierarchical Organization

The computation of Intrinsic Ignition is based on network theory and binarization techniques (Methods and Figure 2a). An example of the raster plot generated for each subject is shown in Figure 2b. In this case, the raster plot is calculated for one subject in six conditions. The ignition capability generates two measures: Intrinsic Ignition and Ignition Variability.

**Figure 2.**
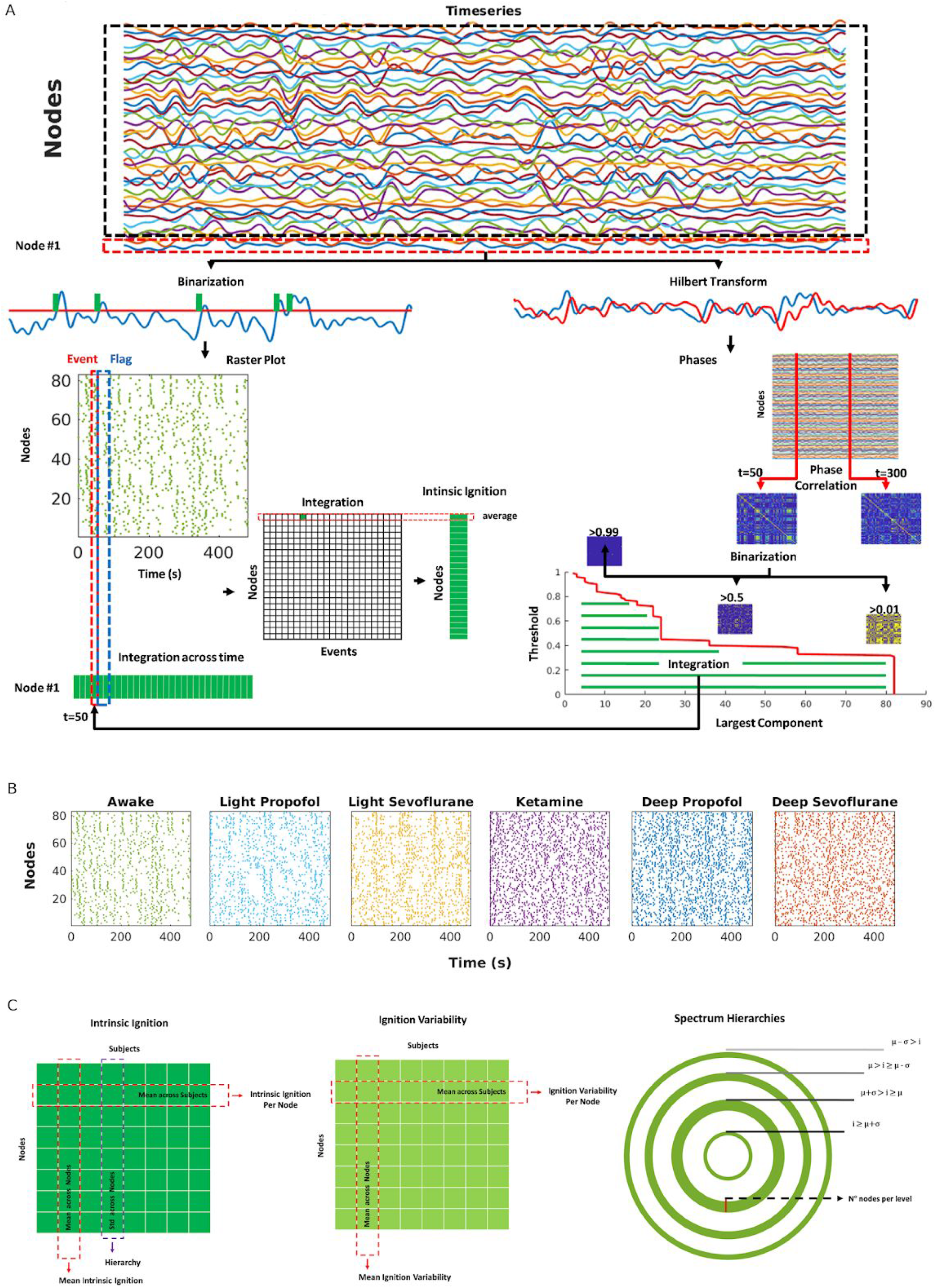
Intrinsic Ignition Measures. **A)** From a binarization (Methods), a raster plot is generated for all nodes. In parallel, a Hilbert transform is applied to each time series, from which a phase correlation or pairwise phase synchronization between regions is defined. For each matrix generated (e.g, *t* = 50 and *t* = 300) another binarization process is applied. With the remaining connections, the largest component is computed as the level of integration given by the length of the connected component of that undirected graph. It creates a curve (lower figure) and the area below the curve (green) corresponds to the integration value for that node at that time-point. Running the same procedure for all time-points results in a table of integration across time for each node. Finally, for each event, the total integration is computed as the average across the time window, the event and a flag of 4TR, building a matrix of integration across events. The average integration across events corresponds to the **Intrinsic Ignition**, while the standard deviation integration across the same events is the **Ignition Variability**. **B)** Example of raster plot for one subject in six conditions, 82 nodes. **C)** The previous procedure is repeated through subjects and conditions, creating a data matrix *D_ij_* for each condition, such that *i* corresponds to the nodes and *j* to the subjects. The mean across subjects (*j*) is the **Intrinsic Ignition per node** and **Ignition Variability per node** respectively. The mean across nodes (*i*) defines the **Mean Intrinsic Ignition** and the **Mean Ignition Variability**. The standard deviation of the Mean Intrinsic Ignition (intrinsic ignition across nodes) returns a quantification value of the shape of the Intrinsic Ignition curve, which is defined as the **Hierarchy**. Ultimately, a **Spectrum Hierarchy** is a circle plot with different levels. Each level corresponds to a threshold of the Intrinsic Ignition and/or Ignition Variability curve (see Methods and Supplementary Figure 2). The thickness of each level line is the number of nodes on that level (e.g. red line marks the thickness of level two).

Plotting the Intrinsic Ignition per node values from highest to lower creates a sorted curve. The shape of that curve informs about the possible types of hierarchical organization in a network. Qualitatively, the Intrinsic Ignition per node curve for the awake condition seems to correspond to a graded non-uniform hierarchy, as previously reported (Deco & Kringelbach, 2017; Deco et al., 2017). While the anesthesia curves transit from graded non-uniform hierarchy to less pronounced curve slope, suggesting spatial modifications towards weak non-hierarchies (Figure 3a). However, a zoom on these curves indicates that the graded non-uniform nature is maintained, and what changes are the degrees of this non-uniformity (Supplementary Figure 2). The higher values of Intrinsic Ignition are found in the awake condition, followed by light sevoflurane and light propofol, and later deep propofol, deep sevoflurane, and ketamine anesthesia. Ketamine anesthesia has a similar effect than deep propofol and deep sevoflurane anesthesia. The curves seem to differentiate at least two groups: awake and anesthesia.

**Figure 3.**
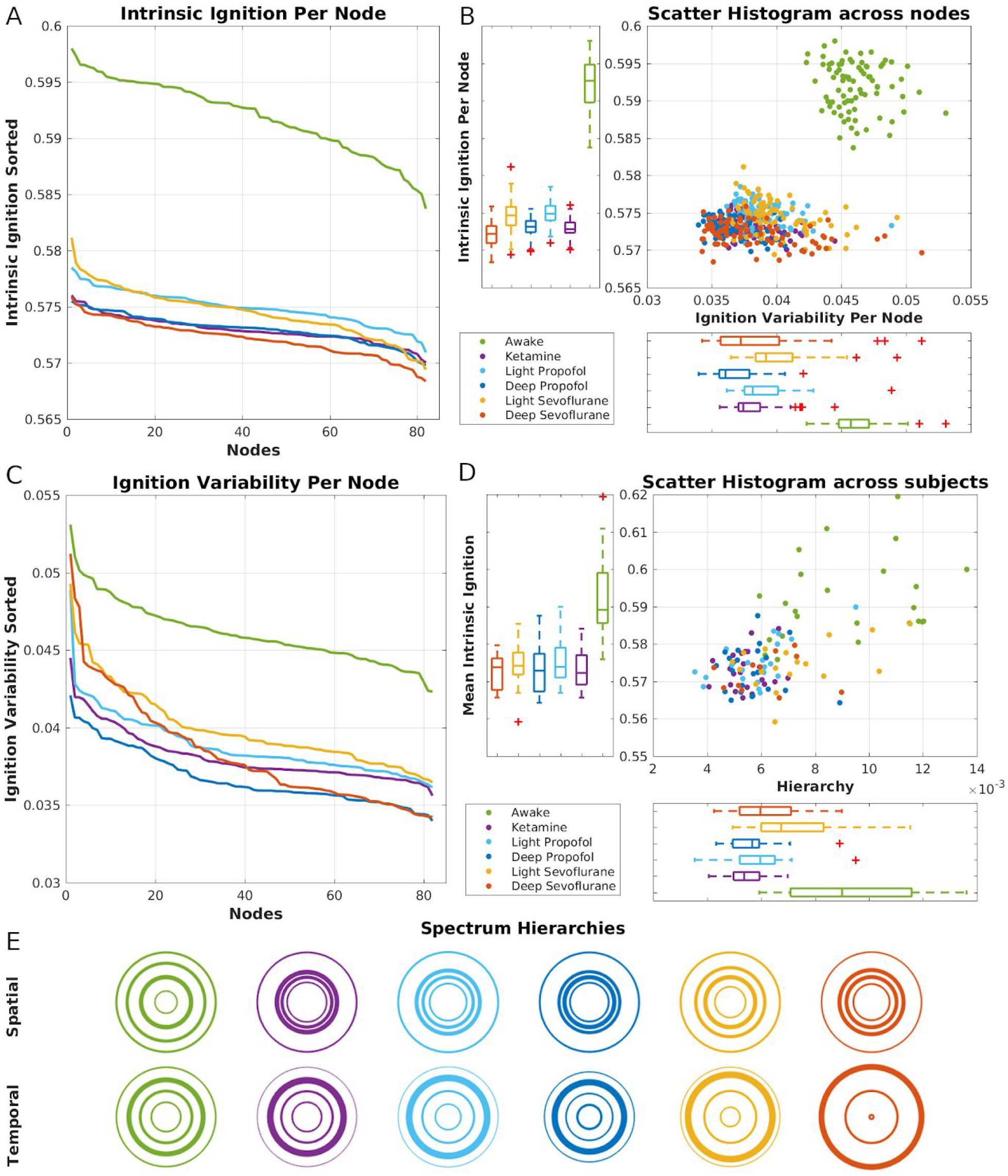
Intrinsic Ignition reveals hierarchical disruption. **A)** The shape of ignition curves changes slightly across conditions, suggesting spatial modifications towards weak non-hierarchies. Nodes are indexed in descendent order. **B)** Scattering plot shows two different groups, awake and anesthesia. Upper right, awake group is significantly different than other conditions (Kolmogorov–Smirnov test p<0.001, awake CI [0.5926 0.5912], ketamine CI [0.5732 0.5727], light propofol CI [0.5752 0.5746], deep propofol CI [0.5733 0.5727], light sevoflurane CI [0.5750 0.5740], deep sevoflurane CI [0.5725 0.5718]). Bottom left, awake is again differentiated from the others conditions (Kolmogorov–Smirnov test p<0.001, awake CI [0.0465 0.0456], ketamine CI [0.0384 0.0377], light propofol CI [0.0392 0.0384], deep propofol CI [0.0371 0.0363], light sevoflurane CI [0.0405 0.0394], deep sevoflurane CI [0.0391 0.0376]). **C)** Ignition Variability curves suggest more intricate ways to disrupt the temporal organization. **D)** Scatter histogram. Awake condition is significantly different than sedations (upper right, Kolmogorov–Smirnov test p<0.001, awake CI [0.5963 0.5875], ketamine CI [0.5750 0.5708], light propofol CI [0.5774 0.5724], deep propofol CI [0.5756 0.5705], light sevoflurane CI [0.5773 0.5716], deep sevoflurane CI [0.5750 0.5693]), as well as across the Hierarchy (bottom left, Kolmogorov–Smirnov test p<0.005, awake CI [0.0102 0.0083], ketamine CI [0.0058 0.0052], light propofol CI [0.0064 0.0053], deep propofol CI [0.0061 0.0052], deep sevoflurane CI [0.0070 0.0053]), with the exception of light sevoflurane (p=0.08, light sevoflurane CI [0.0081 0.0064]). **E)** Spectrum Hierarchies, spatial and temporal. Different changes suggest that disturbing one of the two dimensions of the hierarchical organization may be enough to cause loss of consciousness. For each Box Plot, the bottom and top edges of the box indicate the 25th and 75th percentiles, respectively. The whiskers extend the extreme values without outliers, while outliers are marked as a red cross. The center indicates the median.

Taking both the Intrinsic Ignition value and the Ignition Variability generate a scatter plot with one point per node (Figure 3b). Two clusters are now clearly separated; one corresponds to the awake condition (green dots) and the others to sedation conditions. The box plots, on the right side, shows the distribution of Intrinsic Ignition values across nodes in each condition. Awake values are significantly higher than other anesthetic conditions (Kolmogorov–Smirnov test p<0.001, CI reported in captions Figure 3b), supporting the idea of maximal hierarchical organization. Values for ketamine anesthesia differentiate from light propofol, light and deep sevoflurane (p<0.001) but not deep propofol anesthesia (p>0.01). Light propofol anesthesia presents slightly higher values of Intrinsic Ignition per node, becoming statistically differentiated from deep propofol and deep sevoflurane anesthesia (p<0.001) but not from light sevoflurane anesthesia (p>0.01) which is the third-highest value after the awake condition and light propofol sedation. Intrinsic Ignition in deep propofol anesthesia is significantly lower than in light sevoflurane anesthesia (p<0.001) and slightly higher than in deep sevoflurane anesthesia (p=0.0013). Finally, light and deep sevoflurane anesthesia also present statistical differences as shown in the box plot. All the Intrinsic Ignition per node values across different subjects (full distribution) lead to similar conclusions (Supplementary Figure 3a). Effect size analyses, performed to quantify these differences, also support these findings (Supplementary Figure 4a and Supplementary Table 1). These results support the idea that the spatial dimension regarding the hierarchical organization is disrupted differently among conditions. This disruption is classified in at least three clusters: Awake, Light and Deep sedation effects.

In terms of how these hierarchical disruptions affect the temporal dimension, the distribution of Ignition Variability per nodes across conditions is also plotted in the lower part of Figure 3b. Statistical tests suggest that the Ignition Variability values are more sensitive to the effect of each anesthetic. All the conditions are differentiated (Kolmogorov–Smirnov test p<0.001) between them, with the exception of deep propofol and deep sevoflurane anesthesia (p=0.02). The awake condition, once again, presents the highest Ignition Variability per node value, followed by light sevoflurane, light propofol, ketamine, deep sevoflurane and deep propofol anesthesia (Figure 3b, c, and Supplementary Figure 3b). The effect size analysis is also in agreement with these results (Supplementary Figure 4b and Supplementary Table 1). It indicates that disruption effects are bigger among the temporal dimension of hierarchical organization, especially at the moment of differentiating conditions. To look for these effects, the Ignition Variability curves are plotted in Figure 3c (values are sorted from highest to lowest). The shapes of the Ignition Variability curves seem to capture more complex relations. One example is the case of deep sevoflurane anesthesia, higher values are close to the values of light sevoflurane, while lower values are near deep propofol anesthesia. These results suggest more intricate ways to disrupt the temporal organization than the observed spatial network hierarchy.

One form to quantify the hierarchical disruption is by using the standard deviation of the Intrinsic Ignition curve across nodes, generating one value for each subject (Method and Figure 2c). This, together with the mean across nodes, defined as the Mean Intrinsic Ignition value per each subject, can characterize the spatial hierarchy for each subject. Using both values per subjects, the mean of Intrinsic Ignition and Hierarchy produces another scatter plot in Figure 3d. The clusters are not as evident as before, instead, the points tendency shows a correlation between Mean Intrinsic Ignition and Hierarchy values (Pearson coefficient 0.6, CI [0.47 0.70]). Furthermore, in terms of the Mean Intrinsic Ignition, the awake condition is significantly different from the anesthetics (upper box plot, Kolmogorov–Smirnov test p<0.001, CI reported in captions Figure 3d), while no anesthetics are differentiated between them (Kolmogorov–Smirnov test, p>0.01). Similar analyses on the mean of Ignition Variability are found in Supplementary Figure 5. In terms of hierarchy, the awake condition is also differentiated (bottom box plot, Kolmogorov–Smirnov test p<0.001, CI reported in captions Figure 3d), with the exception of light sevoflurane anesthesia (p=0.08, light sevoflurane). Hence, light sevoflurane anesthesia is the only anesthetic condition which seems to be distinguished from the others (p<0.05), with the exception of deep sevoflurane anesthesia (p=0.39).

Another form to characterize the hierarchical structure is the spectrum hierarchies of each condition (Figure 3e and Supplementary Figure 2). This is a plot that takes the mean and standard deviation of the Intrinsic Ignition per node curve (Spatial, Figure 3e upper) and Ignition Variability per node curve (Temporal, Figure 3e bottom) to describe levels and number of areas for each level. The spatial graphs show how the spectrum changes from a graded non-uniform in the awake condition to a different type of graded and non-uniformity under anesthesia. Among anesthetics, each spectrum presents non-evident visual changes from one spectrum to the other, with only slight changes on light and deep sevoflurane anesthesia (as Hierarchy tests confirmed above), suggesting again that the spatial dimension of the hierarchical organization does not change dramatically across anesthetics. Quite intriguing, if the temporal spectrum is now observed among conditions, no huge differences are perceived from awake state, ketamine, and light propofol anesthesia, but some differences appear in comparison of the awake state, deep propofol, light, and deep sevoflurane anesthesia. This seems to be the opposite tendency from the spatial spectrum. These two different types of changes on the spectrum hierarchies support the idea that disturbing one of the two dimensions (spatial or temporal) may be enough to cause loss of consciousness. It suggests more complex structural qualitative differences concerning spectrum hierarchies that need to be solved in terms of the local number of nodes per level. For example, different regions seem to take different dynamical roles across different conditions, in order to preserve part of the hierarchical organization (Supplementary Tables 2 and 3). Everything together indicates that there is a hierarchical disruption between the awake condition and the anesthetics conditions, while under anesthesia the global values of hierarchy seem to correspond to a similar organization. To understand what is changing and what is not, a local analysis of the differences among nodes is needed.

### 2.3 Local hierarchical Organization

Following the results in Figure 3, each value of Intrinsic Ignition and Ignition Variability per node was plotted in Figures 4 and 5 to show how local differences evolve across conditions. Moreover, a local analysis is performed to isolate regions that may have a bigger impact on the global changes of spatial and temporal hierarchy disruption (Figure 6). To find those regions, the global tendency was defined in terms of three logical propositions from previous analyses (Methods, Figure 2). This global tendency corresponds to higher values of Intrinsic Ignition and Ignition Variability per node during awake (*nodes*≥μ), middle values in light sedation (μ + σ≥*nodes*) and lower levels during deep sedation (μ≥*nodes*). Each node was classified in terms of the spectrum hierarchies levels (Supplementary Tables 2 and 3). Nodes satisfying the logical proposition per condition were defined as a potential area to follow/drive the global tendency observed (red indexes in Supplementary Tables 2 and 3). To visualize the areas and their changes, all values were rescaled/normalized, taking the lower value as zero and the highest as 1 (other normalizations present similar visualizations, data not shown). In Figures 4 and 5, results for Intrinsic Ignition per node and Ignition Variability are plotted respectively. According to this intuitive classification/level method, regions that follow the global tendency are distinguished from others. In the case of the Intrinsic Ignition, these nodes are the right subgenual cingulate cortex, the right posterior cingulate cortex, the right inferior parietal cortex, the right intraparietal cortex, the right frontal eye field, the left parahippocampal cortex, the left subgenual cingulate cortex, the left primary somatosensory cortex, the left intraparietal cortex and the left superior parietal cortex. For Intrinsic Variability, the regions are the right temporal polar, the right central temporal cortex, the right subgenual cingulate cortex, and the left dorsomedial prefrontal cortex. In Figures 4 and 5, these regions are signaled by red arrows (names indexes reported in Supplementary Table 4).

**Figure 4.**
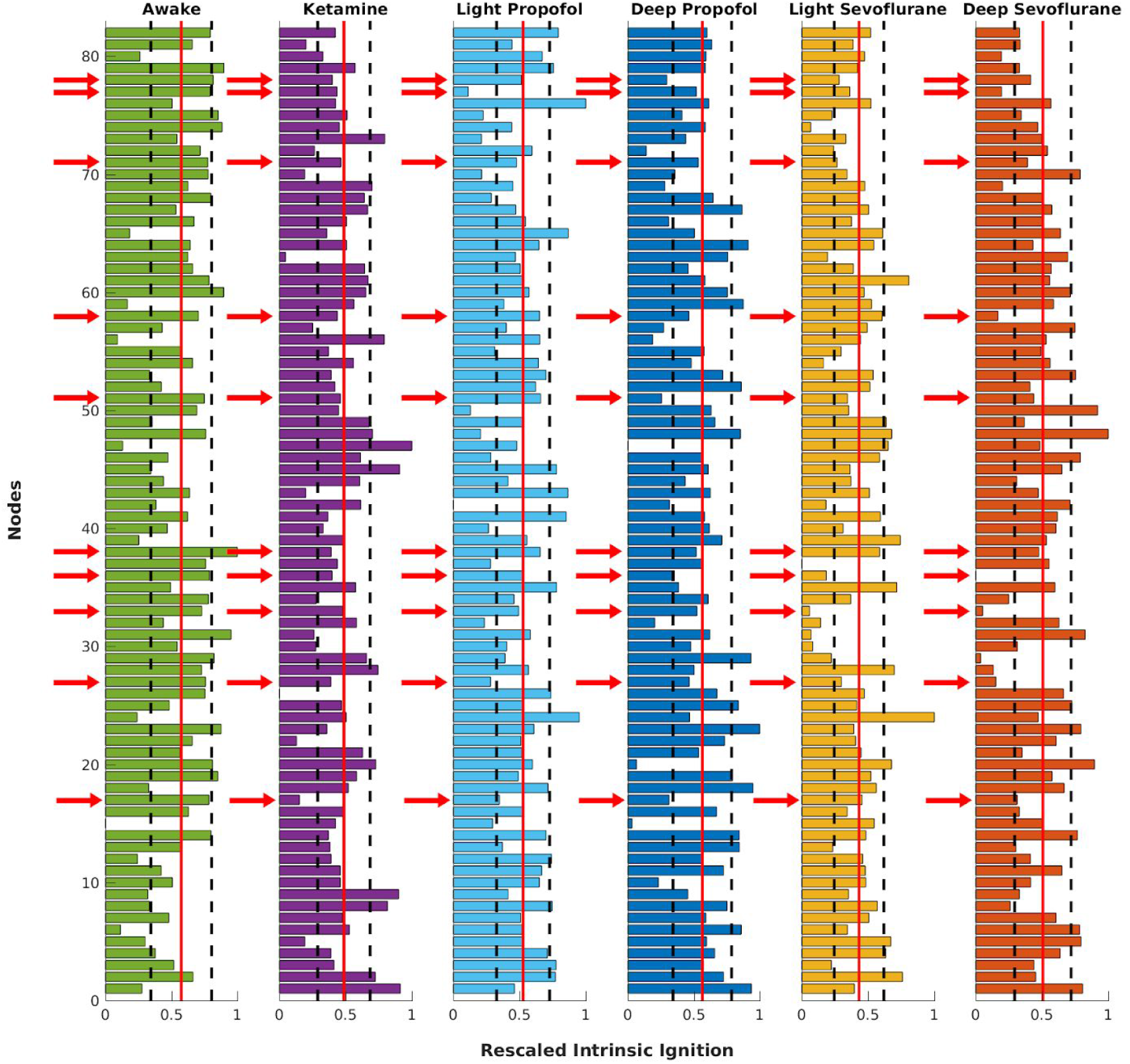
Local tendency of Intrinsic Ignition among conditions. The absolute value of Intrinsic Ignition per node was rescaled from zero (minimal value) to one (max value), allowing a visual comparison across nodes and conditions. The red vertical line corresponds to the mean value of the distribution (μ) and black vertical lines are the limits given by μ±σ, with σ the standard deviation for each condition. The global tendency was translated into three logical propositions: *nodes*≥μfor awake, μ + σ≥*nodes* for light conditions and μ≥*nodes* for deep conditions. Only 10 regions (Subgenual cingulate cortex right, Posterior cingulate cortex right, Inferior parietal cortex right, Intraparietal cortex right, Frontal eye field right, Parahippocampal cortex left, Subgenual cingulate cortex left, Primary somatosensory cortex left, Intraparietal cortex left, Superior parietal cortex left) satisfied the three propositions simultaneously (Supplementary Tables 3, 4 and 5), becoming candidates for areas which follow the global tendency of Intrinsic Ignition changes. These regions are signaled by red arrows and correspond to the indexes 17, 27, 33, 36, 38, 51, 58, 71, 77, and 78 respectively (Supplementary Table 9).

**Figure 5.**
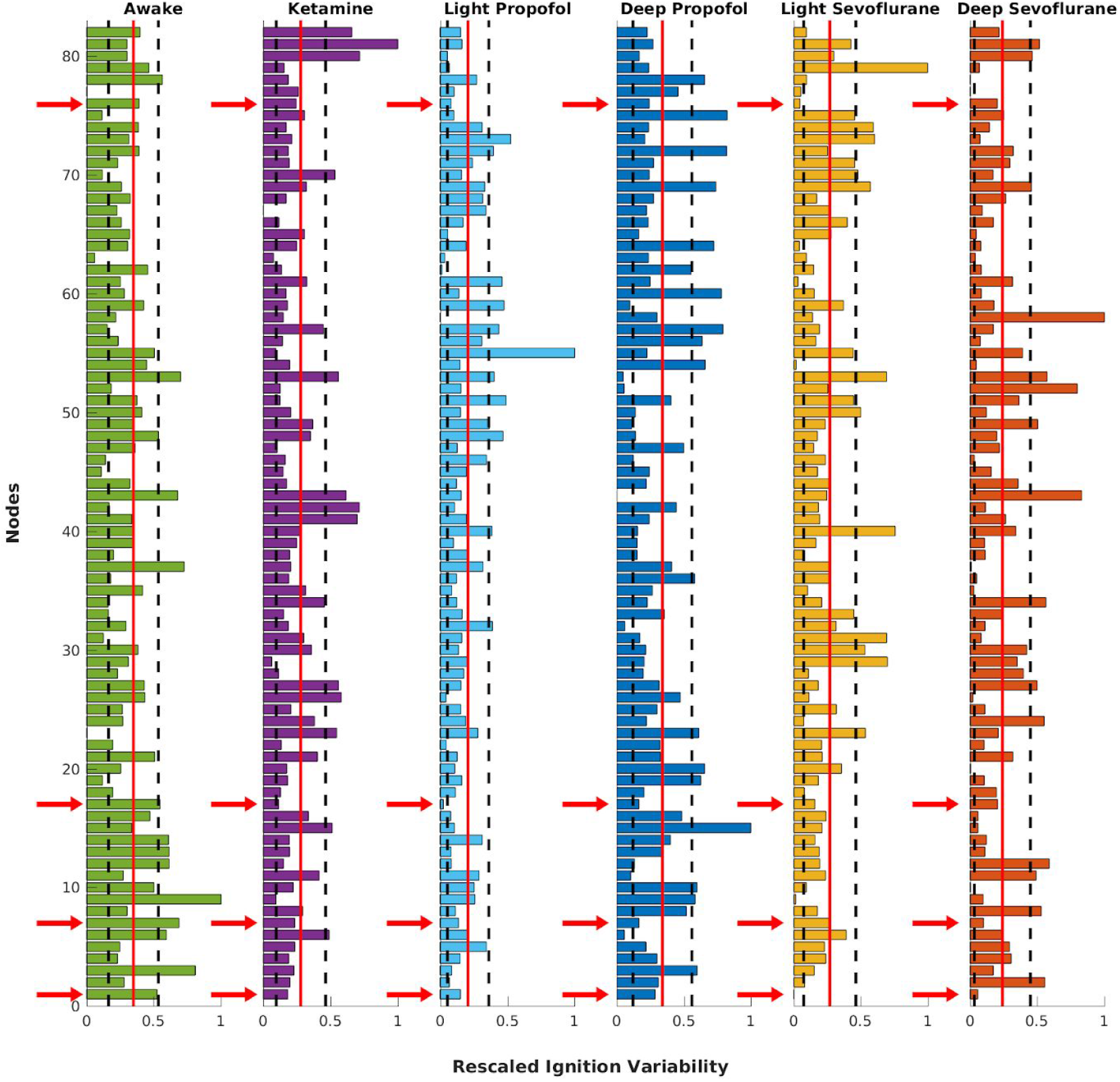
Local tendency Ignition Variability among conditions. The red vertical line corresponds to the mean value of the distribution (μ) and black vertical lines are the limits given by μ± σ, with σ the standard deviation for each condition. The global tendency was translated into three logical propositions: *nodes*≥μ for awake, μ + σ≥*nodes* for light conditions and μ≥*nodes* for deep conditions. Only 4 regions (Tempolar polar right, Central temporal cortex right, Subgenual cingulate cortex right, Dorsomedial prefrontal cortex left) survived the three propositions simultaneously (Supplementary Tables 6, 7 and 8), becoming candidates for areas which follow the global tendency of ignition variability changes. These regions are signaled by red arrows and correspond to the indexes 1, 7, 17, and 76 respectively (Supplementary Table 9).

**Figure 6.**
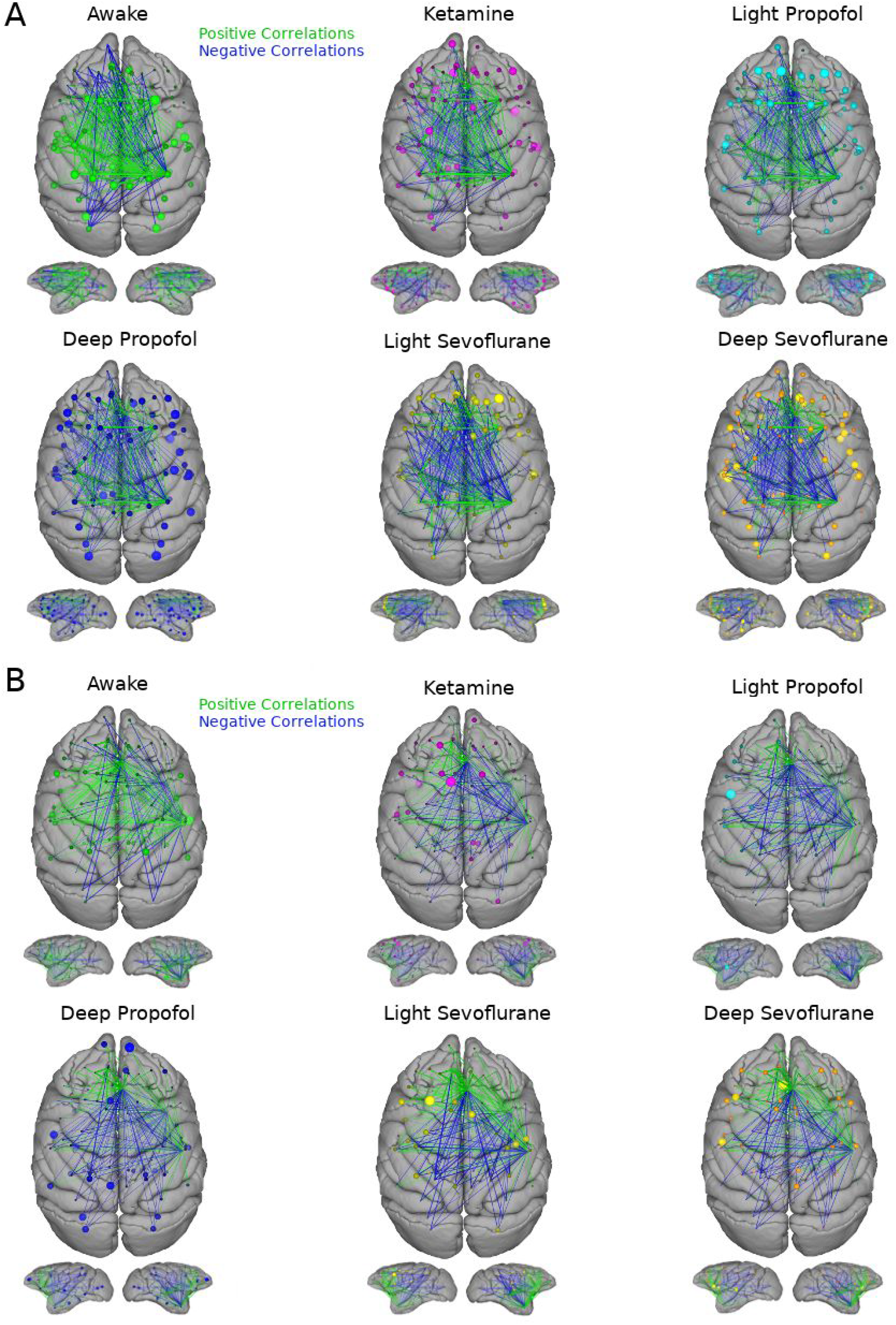
Global and local aspects of the brain functional network. A graphical way to observe how local and global disruption are intrinsically connected but probably differently driven under dissimilar anesthetics. **A)** Intrinsic Ignition driven network. For each condition and node, the values of Intrinsic Ignition are plotted as the size of the node, while the links correspond to the functional correlation only for the nodes identified as relevant from Figure 4. Changes among both sizes as well as the number of links at different anesthetics are observed, suggesting interdependency between the local (nodes) and global (network) changes. **B)** Similar plots for Ignition Variability driven network with the functional correlation only for the relevant nodes from Figure 5. In this case, changes among node sizes seem more relevant than among the number of links, suggesting that Ignition Variability may be more sensitive to these local (node) changes.

Supplementary analyses were performed using the median instead of the mean as a cut off on the logical propositions (plots available by request). It slightly changes the results for Intrinsic Ignition. In the case of Ignition Variability, using the median instead of the mean gives no results. Finally, to ensure consistency, the same analyses were performed subject by subject (see Methods), generating a histogram of occurrences. Nodes that appear to satisfy the logical propositions per conditions above the 60% of the time were identified. These regions are the right subgenual cingulate cortex and the right intraparietal cortex for Intrinsic Ignition (Supplementary Figure 6b), and only the right central temporal cortex, for Ignition Variability (Supplementary Figure 6b).

## 3. Discussion

In this study, we analyzed resting-state fMRI acquired from non-human primates in the awake state and during anesthesia-induced loss of consciousness using distinct pharmacological agents. By applying Intrinsic Ignition measurements, we demonstrate that loss of consciousness is paralleled by a disruption of brain hierarchy, making it both a new signature of consciousness and consciousness loss.

At the neuronal level, anesthetic agents act through different molecular pathways (Hudetz & Mashour, 2016), such as GABA receptors, NMDA receptors and K+ channels (Franks, 2008) to induce loss of consciousness. At the system level, anesthetics strongly affect brain networks that are involved in arousal and awareness. Propofol (V. Bonhomme et al., 2001), sevoflurane (Kaisti et al., 2003) and xenon (Laitio et al., 2007) inhibit midbrain areas, associated with the ascending reticular system, the thalamus and cortical areas such as the precuneus, posterior cingulate cortex, and the prefrontal cortex. Moreover, studies with ketamine, propofol, and sevoflurane (Lee et al., 2013; Uhrig et al., 2018) reported disruption of frontoparietal activity despite their distinct neurophysiology. Ketamine and sevoflurane have opposite effects on thalamocortical connectivity. Under ketamine, an N-methyl-D-aspartate receptors antagonist (Anis, Berry, Burton, & Lodge, 1983), thalamocortical functional correlations seems to be preserved (Bonhomme et al., 2016), while sevoflurane, a γ-aminobutyric acid receptor type A agonist and N-methyl-D-aspartate antagonist (Wu, Harata, & Akaike, 1996), induces a decrease in thalamocortical functional correlations, but preserving functional correlations between thalamus and sensory cortex (Huang et al., 2014; Ranft et al., 2016). Propofol, targeting γ-aminobutyric acid-mediated neurotransmission (Peduto, Concas, Santoro, Biggio, & Gessa, 1991), induces disconnections in thalamocortical circuits. Under anesthesia, the brain patterns are not uniform, some areas seem more deactivated than others, while most anesthetics, with the exception of ketamine (Langsjo et al., 2005), cause a global reduction of cerebral blood flow (Franks, 2008). Taking this evidence, finding a common neurophysiological pathway for this different anesthetics is challenging, but might involve the disruption of thalamocortical connectivity and/or frontoparietal connectivity.

Our results indicate that Intrinsic Ignition can discriminate between the awake condition and different anesthetics, as well as between different levels of sedation (i.e. light versus deep anesthesia). Moreover, higher values of ignition are associated with richer brain dynamics, while lower values relate to structural driven dynamics, as shown in previous reports (Barttfeld et al., 2015; Uhrig et al., 2018; Demertzi et al., 2019). Our results (Figure 1) are in line with these previous studies, and additionally indicate that the spatial and temporal dimensions of hierarchical organization change under anesthesia, while keeping different types of the same graded non-uniform hierarchy through all conditions. Different regions occupy different levels on those hierarchies, indicating that different anesthetic paths may indeed act differently, however, generating the same hierarchical disruption. Everything together supports the hierarchical breakdown hypothesis and places the local and global disruption of the hierarchy as a possible common mechanism to reconcile seemingly different types of loss of consciousness.

Reports from NREM sleep also provide evidence for a partial breakdown of the hierarchical organization of large scale networks (Boly, Perlbarg, et al. 2012). In order to quantify these disruptions, Intrinsic Ignition (Figure 3), was applied in two hierarchical dimensions: spatial and temporal. The awake condition in monkeys presents maximal hierarchical organization (Figure 3), while different concentrations of anesthetics range from middle to lower values of Intrinsic Ignition, under light and deep anesthesia (Figure 3b and d). Changes observed on the spatial dimension (Intrinsic Ignition per node), recognized three groups: awake, light (propofol and sevoflurane) and deep (propofol, sevoflurane, and ketamine). On the other hand, changes observed on the temporal dimension (Intrinsic Variability per node) seems more sensitive, distinguishing among all the conditions (Figure 3b and c) and suggesting more complex hierarchical disruptions across time. Interestingly, light conditions presented middle values on both Intrinsic Ignition (spatial) and Ignition Variability (temporal), while ketamine was closer to deep conditions under spatial dimension but near to light conditions when compared across the temporal aspect. This suggests that the previous evidence on the dynamical disruptions is related to more complex multidimensional disruptions given by at least two dimensions: structural organization (spatial) and dynamical organization (temporal). It supports the idea that spatial and temporal hierarchies are affected differently among conditions.

The degree of Hierarchy seems disturbed among all the different sedation (with the exception of light sevoflurane). It may imply that hierarchical disruptions are indeed more complexly related to possible local mechanisms. The Hierarchy quantifier recognizes that light and deep sevoflurane anesthesia might be affecting similarly but in different degrees the hierarchical organization, in turn that estimates that light sevoflurane anesthesia may have weaker effects on that disruption and still closer to an awake organization. If this interpretation is right, ketamine, light, and deep propofol anesthesia would break similarly the hierarchical organization, since they present similar distributions in terms of Hierarchy. Moreover, spectrum hierarchies reveal similar spatial disruptions between ketamine, light, and deep propofol anesthesia (Figure 3e upper) and partially similar ones in the temporal dimension (Figure 3e lower). It indicates that different mechanisms disrupt one or another aspect of hierarchical organizations: the spatial dimension conveys the impression that some anesthetics are distinguished but not others, while in the temporal dimension it happens among different anesthetics. Plot curves (Figure 3a and c) and spectrum plots (Figure 3e, Supplementary Figure 2) show that graded non-uniform type of hierarchy is maintained across conditions. Disturbing one of the two dimensions may be enough to cause loss of consciousness and different anesthetics target differently these two aspects of the neural organization: ketamine and propofol would target spatial aspects, while sevoflurane would disturb temporal aspects of that configuration.

To deal with these structural differences, local analyses were performed. According to our results, organizational disruptions cannot be reduced to only global effects but also local differences (Figures 4, 5, 6 and Supplementary Tables 2 and 3). Globally, some anesthetics may act similarly in terms of concentration, such as light propofol and light sevoflurane anesthesia, however, locally, different anesthetics may also present differences given mainly in terms of Ignition Variability curves (Figure 3). This is clearly noticed from the different distributions of values for Intrinsic Ignition and Ignition Variability per node presented in Figures 4 and 5. Once observed which areas followed the global tendency as candidates for driving these changes, Subgenual cingulate cortex and Intraparietal cortex presented a consistent occurrence in terms of Intrinsic Ignition, and Central temporal cortex in terms of Ignition Variability. Additionally, Posterior cingulate cortex, Inferior and Superior parietal cortex, Parahippocampal cortex, and somatosensory cortex among other prefrontal regions appeared as relevant areas for spatial aspects of ignition. Most of these regions have been previously associated with GNW (Uhrig et al., 2014; Uhrig et al., 2016, 2018). In terms of the temporal component, Temporal polar cortex, Subgenual cingulate cortex, and Dorsomedial prefrontal cortex also seem to play a role in the global disruptions among the six conditions.

These local findings are in line with previous reports. For instance, studies on anesthesia and early stages of sleep have identified that the precuneus and lateral temporoparietal components of DMN persisted under anesthesia (Vincent et al., 2007), but the connectivity of the posterior cingulate cortex (PCC) is reduced during sedation (Kaisti et al., 2002; Greicius et al., 2008; Uhrig et al., 2018). Moreover, studies using light propofol point out changes among PCC connectivity with other areas, such as somatomotor cortex, the anterior thalamic nuclei and the reticular activating system (Stamatakis, Adapa, Absalom, & Menon, 2010). According to our results, the PCC is following the global tendency among conditions and therefore, is a candidate for globally driving the hierarchical changes observed. An fMRI meta-analysis of resting-state activity in disorders of consciousness concluded that a reduction of activity in midline cortical (PCC, precuneus, medial temporal lobe, middle frontal lobe) and subcortical sites (bilateral medial dorsal nuclei of the thalamus) is associated with conscious impairments (Hannawi, Lindquist, Caffo, Sair, & Stevens, 2015), with a more pronounced reduction in the vegetative state than in the minimally conscious state. Moreover, medial parietal cortex, PCC and precuneus are the first regions to reactivate when patients recover (Laureys, Boly, & Maquet, 2006). Due to the current parcellation, our method cannot target all these areas, nevertheless, according to intrinsic ignition, some of them may be related to the global changes observed.

Our results, however, are not free of limitations. The data analyzed was with a parcellation of only 82 cortical regions of interest (CoComac) and therefore does not allow to infer the disruption of the whole brain hierarchical organization. A subcortical parcellation and better definition for SC (although not mostly used here) are desirable for these effects (Kennedy, Knoblauch, & Toroczkai, 2013). The preprocessing pipeline can be improved (Tasserie et al., 2019) in order to avoid the extra cleaning procedure. Although our results indicate similar global hierarchical disruptions as a common mechanism driven by locally different re-organizations, these results need modelling and simulations to give a full answer about the casual driving disruptions. For example in (Chaudhuri, Knoblauch, Gariel, Kennedy, & Wang, 2015) and (Joglekar, Mejias, Yang, & Wang, 2018) the ignition capability was explored as an inter-areal balanced amplification signal through large scale circuits, supporting ignition models of consciousness (Joglekar et al., 2018). Moreover, temporal hierarchies naturally emerged from the heterogeneity of local networks (Chaudhuri et al., 2015), with slower prefrontal and temporal regions having a strong impact on global brain dynamic. Therefore, in order to link intrinsic ignition and mechanistic models, large scale models (Breakspear, 2017) are expected as future steps to give light on part of the neuronal mechanisms involved. It may help to connect our results with other studies on sleep (Jobst et al., 2017) and disorders of consciousness, as well as the simulation and exploration of manipulated brain states using deep brain stimulation (Saenger et al., 2017).

## 4. Conclusion

In conclusion, the global values of hierarchical organization indicate similar global organization (disruptions) under anesthesia, while local analyses on which areas habit hierarchical levels inform on the different ways that anesthetics affect spatial and temporal aspects of that organization. Our study provides a common brain signature of anesthesia-induced loss of consciousness beyond molecular pharmacology, also called “common anesthetic endpoint” (Hudetz & Mashour 2016). This is in line with the idea that disruptions in long-distance network dynamics are a common signature of anesthesia-induced loss of consciousness, but adding the breakdown of hierarchical organization and its two dimensions, space and time. The hierarchical organization is characterized by internal ignition activity, reconciling the observed common global changes (hierarchies) with different local changes (ignition power by node).

Our local results suggest that areas proposed by GNW, such as fronto-parieto-cingular networks, which underpin conscious access (Dehaene & Changeux, 2011), and regions considered by IIT as the parietal-posterior cortical zones (supporting subjective experience (Siclari et al., 2017)), are both participating in changes of hierarchical organization. These hierarchical changes find their common ground in cingulate and parietal regions. It may imply, that under the hierarchical hypothesis, both theories are complementary to each other, and approaching compatible aspects of the same conscious phenomenon (Aru, Bachmann, Singer, & Melloni, 2012; Block, 2005; Dehaene et al., 2014; Tagliazucchi, 2017; Mashour et al., 2020). This solves in part the requirement of a desirable common and global mechanism to explain how brain dynamics are similarly affected under anesthetics, at the same time than recovering the specificity of affecting and modulating correlations and couplings of brain regions.

## 5. Methods

### 5.1 Animals

The acquisition of this data set is previously reported in (Barttfeld et al., 2015) and (Uhrig et al., 2018, http://links.lww.com/ALN/B756). Five rhesus macaques were included for analyses (*Macaca mulatta*, one male, monkey J, and four females, monkeys A, K, Ki, and R, 5-8 kg, 8-12 yr of age), in a total of six different arousal conditions: Awake state, ketamine, light propofol, deep propofol, light sevoflurane, and deep sevoflurane anesthesia. Three monkeys were used for each condition: Awake (monkeys A, K, and J), Ketamine (monkeys K, R and Ki), Propofol (monkeys K, R, and J), Sevoflurane (monkeys Ki, R, and J). All procedures are in agreement with the European Convention for the Protection of Vertebrate Animals used for Experimental and Other Scientific Purposes (Directive 2010/63/EU) and the National Institutes of Health’s Guide for the Care and Use of Laboratory Animals. Animal studies were approved by the institutional Ethical Committee (Commissariat à l’Énergie atomique et aux Énergies alternatives; Fontenay aux Roses, France; protocols CETEA #10-003 and 12-086).

### 5.2 Anesthesia Protocols

The anesthesia protocol is thoroughly described in previous studies (Barttfeld et al., 2015; Uhrig et al., 2018). Monkeys received anesthesia either with ketamine (Uhrig et al., 2018), propofol (Barttfeld et al., 2015) or sevoflurane (Uhrig et al., 2018), with two different levels of anesthesia depth for propofol and sevoflurane anesthesia (Light and Deep). These levels were defined according to the monkey sedation scale, based on spontaneous movements and the response to external stimuli (presentation, shaking or prodding, toe pinch), and corneal reflex. For each scanning session, the clinical score was determined at the beginning and end of each scanning session, together with continuous electroencephalography monitoring (Uhrig et al., 2016).

During ketamine, deep propofol anesthesia, and deep sevoflurane anesthesia, monkeys stopped responding to all stimuli, reaching a state of general anesthesia. Monkeys were intubated and ventilated as previously described (Barttfeld et al., 2015; Uhrig et al., 2018). Heart rate, noninvasive blood pressure, oxygen saturation, respiratory rate, end-tidal carbon dioxide, and cutaneous temperature were monitored (Maglife, Schiller, France) and recorded online (Schiller).

Ketamine was applied by intramuscular injection (20 mg/kg; Virbac, France) for induction of anesthesia, followed by a continuous intravenous infusion of ketamine (15 to 16 mg · kg–1 · h–1) to maintain anesthesia. Atropine (0.02 mg/kg intramuscularly; Aguettant, France) was injected 10 min before induction, to reduce salivary and bronchial secretions. For propofol, monkeys were trained to be injected an intravenous propofol bolus (5 to 7.5 mg/kg; Fresenius Kabi, France), followed by a target-controlled infusion (Alaris PK Syringe pump, CareFusion, USA) of propofol (light propofol sedation, 3.7 to 4.0 μg/ml; deep propofol anesthesia, 5.6 to 7.2 μg/ml) based on the “Paedfusor” pharmacokinetic model (Absalom & Kenny, 2005). During sevoflurane anesthesia, monkeys received first an intramuscular injection of ketamine (20 mg/kg; Virbac) for induction, followed by sevoflurane anesthesia (light sevoflurane, sevoflurane inspiratory/expiratory, 2.2/2.1 volume percent; deep sevoflurane, sevoflurane inspiratory/expiratory, 4.4/4.0 volume percent; Abbott, France). Only 80 minutes after the induction the scanning sessions started to get a washout of the initial ketamine injection (Schroeder et al., 2016). To avoid artifacts related to potential movements throughout magnetic resonance imaging acquisition, a muscle-blocking agent was coadministered (cisatracurium, 0.15 mg/kg bolus intravenously, followed by continuous intravenous infusion at a rate of 0.18 mg · kg–1 · h–1; GlaxoSmithKline, France) during the ketamine and light propofol sedation sessions.

### 5.3 Functional Magnetic Resonance Imaging Data Acquisition

For the awake condition, monkeys were implanted with a magnetic resonance–compatible head post and trained to sit in the sphinx position in a primate chair (Uhrig, Dehaene, & Jarraya, 2014). For the awake scanning sessions, monkeys sat inside the dark magnetic resonance imaging scanner without any task and the eye position was monitored at 120 Hz (Iscan Inc., USA). For the anesthesia sessions, animals were positioned in a sphinx position, mechanically ventilated, and their physiologic parameters were monitored. Before each scanning session, a contrast agent, monocrystalline iron oxide nanoparticle (Feraheme, AMAG Pharmaceuticals, USA; 10 mg/kg, intravenous), was injected into the monkey’s saphenous vein (Vanduffel et al., 2001). Monkeys were scanned at rest on a 3-Tesla horizontal scanner (Siemens Tim Trio, Germany) with a single transmit-receive surface coil customized to monkeys. Each functional scan consisted of gradient-echoplanar whole-brain images (repetition time = 2,400 ms; echo time = 20 ms; 1.5-mm3 voxel size; 500 brain volumes per run).

### 5.4 Functional Magnetic Resonance Imaging Preprocessing

A total of 157 functional magnetic imaging runs were acquired (Barttfeld et al., 2015; Uhrig et al., 2018): Awake, 31 runs (monkey A, 4 runs; monkey J, 18 runs; monkey K, 9 runs), Ketamine, 25 runs (monkey K, 8 runs; monkey Ki, 7 runs; monkey R, 10 runs), Light Propofol, 25 runs (monkey J, 2 runs; monkey K, 10 runs; monkey R, 12 runs), Deep Propofol, 31 runs (monkey J, 9 runs; monkey K, 10 runs; monkey R, 12 runs), Light Sevoflurane, 25 runs (monkey J, 5 runs; monkey Ki, 10 runs; monkey R, 10 runs), Deep Sevoflurane anesthesia, 20 runs (monkey J, 2 runs; monkey Ki, 8 runs; monkey R, 11 runs). For details, check the supplementary tables for (Uhrig et al., 2018) (http://links.lww.com/ALN/B756).

Functional images were reoriented, realigned, and rigidly coregistered to the anatomical template of the monkey Montreal Neurologic Institute (Montreal, Canada) space with the use of Python programming language and Oxford Centre Functional Magnetic Resonance Imaging of the Brain Software Library software (United Kingdom, http://www.fmrib.ox.ac.uk/fsl/; accessed February 4, 2018) (Uhrig et al., 2014). From the images, the global signal was regressed out to remove any confounding effect due to physiologic changes (*e.g.*, respiratory or cardiac changes). Voxel time series were filtered with a low-pass (0.05-Hz cutoff) and high-pass (0.0025-Hz cutoff) filters and a zero-phase fast-Fourier notch filter (0.03 Hz) to remove an artifactual pure frequency present in all the data (Barttfeld et al., 2015; Uhrig et al., 2018).

Furthermore, an extra cleaning procedure was performed to ensure the quality of the data after time-series extraction (Supplementary Figure 1). The procedure was based on a visual inspection of the time series for all the nodes, the Fourier transform of each signal, the functional connectivity for each subject and the dynamical connectivity computed with phase correlation. Trials were kept when the row signal did not present signs of artifactual activity, functional connectivity was coherent with the average and dynamical connectivity presented consistent patterns across time.

Finally, a total of 119 runs are analyzed in subsequent sections: Awake state 24 runs, ketamine anesthesia 22 runs, light propofol anesthesia 21 runs, deep propofol anesthesia 23 runs, light sevoflurane anesthesia 18 runs, deep sevoflurane anesthesia 11 runs.

### 5.5 Anatomical Parcellation and Structural Connectivity

Anatomical (structural) data were derived from the CoCoMac 2.0 (Bakker, Wachtler, & Diesmann, 2012) database (cocomac.g-node.org) of axonal tract-tracing studies using the Regional Map parcellation (Kötter & Wanke, 2005). This parcellation comprises 82 cortical ROIs (41 per hemisphere; Supplementary Table 4). Structural (*i.e.*, anatomical) connectivity data are expressed as matrices in which the 82 cortical regions of interest are displayed in x-axis and y-axis. Each cell of the matrix represents the strength of the anatomical connection between any pair of cortical areas. The CoCoMac connectivity matrix classifies the strength of the anatomical connections as weak, moderate, or strong, codified as 1, 2, and 3, respectively (Barttfeld et al., 2015).

### 5.6 Dynamic Analyzes

Functional connectivity matrices (FC) for each condition are first computed for each subject using Pearson correlation and then averaged across subjects. Each FC has 82 cortical regions. The dynamical functional connectivity (dFC) is computed using a sliding window technique (50 TR correlation window and 5 TR sliding size). This method is better for visualization purposes. Correlations between FC and Structural connectivity (SC) for each subject are computed with Pearson correlation and plotted as a violin plot. For violin plots, the shape describes the distribution density, the white dot corresponds to the median, the thick inner line is the first quartile (down), and the third quartile (up). The borders are the upper and lower adjacent values (Hintze & Nelson, 1998). Finally, metastability is computed as the standard deviation of the Kuramoto’s order parameter (synchrony).

### 5.7 Intrinsic Ignition Analyzes

The ignition capability can be defined in terms of its spatial and temporal components, generating two measures: Intrinsic Ignition and Ignition Variability. This procedure generates one value for each node and subject that is later averaged to form the Intrinsic Ignition per node and Ignition Variability per node value. Therefore, Intrinsic Ignition tells us about the spatial diversity of a network, while the Ignition Variability, about the diversity across time.

Intrinsic Ignition is a novel technique based on graph and network theory (Figure 2a). For any node, its inner ignition capability is fully characterized by the Intrinsic Ignition as a measure of its spatial diversity, and its variability as a measure of its diversity across time. To compute both aspects, any continuous signal can be binarized using a threshold θ such that the binary sequence *σ_i_*(*t*) = 1 *if z_i_*(*t*) > θ, crossing the threshold from below, and *σ_i_*(*t*) = 0, otherwise (Tagliazucchi, Balenzuela, Fraiman, & Chialvo, 2012). This simple method generates a raster plot with a discrete sequence of events, which is more efficient in terms of complex computations (see raster plots examples for each condition in Figure 2b).

Moreover, a Hilbert transform is performed to the continuous signal, defining the phases for each time point and node. Using these phases, a phase correlation or pairwise phase synchronization between regions *j* and *k* is defined as *P_jk_*(*t*) = *e*^−3|(φ_*j*_−φ_*k*_|^. For each of these matrices, another binarization process is applied for a given absolute threshold θ between 0 and 1 (scanning the whole range), and therefore the symmetric phase lock matrix *P_jk_*(*t*) can be binarized such as 0 *if P_jk_*(*t*)>θ, 1 otherwise. Then the length of the largest component is computed, generating a curve with this value for each binarized phase lock matrix. The area below the curve is defined as the integration value for that node (ROI) at that time point. Running the same procedure for all time points creates a table of integration across time for each node. For each event defined from the first binarization procedure, and a flag window (commonly 4 TR), the total integration is computed as the average across the delta time defined by the event and the flag. It builds a matrix of integration across events. The average integration across events is defined as the **Intrinsic Ignition**, while the standard deviation integration across the same events corresponds to the **Ignition Variability**.

Intrinsic Ignition and Intrinsic Variability produce one value for each of the 82 nodes. The procedure is repeated through subjects and conditions, creating a data matrix *D_ij_* for each condition such that *i* corresponds to the nodes and *j* to the subjects (Figure 2c). The mean of these values across subjects (*j*) corresponds to the **Intrinsic Ignition per node** and **Ignition Variability per node**, respectively. The mean across nodes (*i*) is defined as the **Mean Intrinsic Ignition** and the **Mean Ignition Variability**. The standard deviation of the Mean Intrinsic Ignition (intrinsic ignition across nodes) returns a quantification value of the shape of the Intrinsic Ignition curve (considering all the values sorted from higher to lower), which here is defined as the **Hierarchy**, a quantifier for each subject.

To complement these analyses, a **Spectrum Hierarchy** plot is specified as a circle plot with different levels. Each level corresponds to a threshold of the Intrinsic Ignition (Spatial) and/or Ignition Variability (Temporal) curve: *i*≥μ + σ; μ + σ> *i*≥μ; μ> *i*≥μ-σ; μ − σ > *i. i* refers to the index node, μ the mean value, and σ the standard deviation of the curve. The distance between level is given by the thickness of the level and the value of μ + σ for level 1, μ for level 2, μ-σ for level 3, μ−σ−*min*(*nodes*) for level 4 (Supplementary Figure 2). The thickness of each level line is the number of nodes on that level; a thicker line means more nodes than thinner lines (e.g. red line in the figure marks the thickness of level two). The uniformity of the spectrum hierarchy for one condition characterizes the ignition curves in terms of the hierarchical organization across nodes (Deco & Kringelbach, 2017).

To explore if some nodes would be locally driving the global changes observed, a local analysis was performed on Intrinsic Ignition and Ignition Variability per node. This test consisted of finding which nodes are following the global tendency measured in the spatial and temporal aspects of ignition. As will be discussed in Results, the global tendency was the highest values of Intrinsic Ignition and Ignition Variability in awake, medium values in light sedation, and lower values in deep conditions. These tendencies were translated to logical propositions in order to find the nodes that satisfied all the propositions across conditions. These propositions were: *nodes*≥μ for awake, μ + σ≥*nodes* for light conditions and μ≥*nodes* for deep conditions. More restricted logical propositions do not produce results. Additionally, instead of the mean (μ), the median was also used as cut off in order to search for consistent results. The number of regions obtained from one or another method changes slightly for Intrinsic Ignition, but there is a clear overlap of regions, confirming part of the consistency expected. Finally, a subject by subject analysis was also performed as a supplementary test. In this case, the same propositions above were run in each subject to later generate a histogram of occurrence. A threshold of 60% of occurrence was imposed to find the regions above the threshold.

### 5.8 Statistical Analyzes

The main statistical test used in this work was the non-parametric Kolmogorov–Smirnov test (unless another test is explicitly stated). It is due to the characteristics of the data and their distributions (Rosner, 2012). Therefore, the independence of measures and conditions is a statistical assumption commonly accepted for monkey data (Uhrig et al., 2018), together with the continuous nature of our measures (Rosner, 2012). Confidence intervals (CI) at 95% were computed as 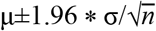. As above, μ is the mean, σ the standard deviation and *n* the length of the data points.

Additionally, effect size analyses were performed to quantify the differences given by statistical tests on Intrinsic Ignition and Ignition Variability per node. The method used was the rank-biserial correlation analysis for independent samples (other effect size techniques such as mean difference, AUROC and Cohen U1 did not present major differences with the results of rank-biserial correlation). In this test, ranks between −1 to +1 correspond to maximal effects and 0 means no effect. To compute a confidence interval for effect size analyzes, 10.000 bootstrapping iterations were performed (more details (Hentschke & Stüttgen, 2011)).

## Acknowledgment

The authors thank Jordy Tasserie for help with data transfer; Morgan Dupont and Wilfried Pianezzola for help with animal experiments; Alexis Amadon, Hauke Kolster, Laurent Laribière, Jérémy Bernard, Eric Giacomini, Michel Luong, Edouard Chazel, and the NeuroSpin magnetic resonance imaging and informatics teams for help with imaging tools; Sébastien Mériaux and Joël Cotton for animal facilities; and Jean-Robert Deverre, for administrative support, Commissariat à l’Énergie atomique et aux Énergies alternatives (CEA /Joliot).

## Contribution

CMS analyzed the data, theoretical design, visualization, and figures, wrote original draft and manuscript edition, LU acquisition data and manuscript edition, MK analytical tools and manuscript edition, BJ acquisition data, experimental design and manuscript edition, GD analytical tools, theoretical design, and manuscript edition. BJ and GD contribute equally to this work.

