## Supplementary Figures for "Hierarchical disruption in the cortex of anesthetized monkeys as a new signature of consciousness loss"

### I. Supplementary Figures

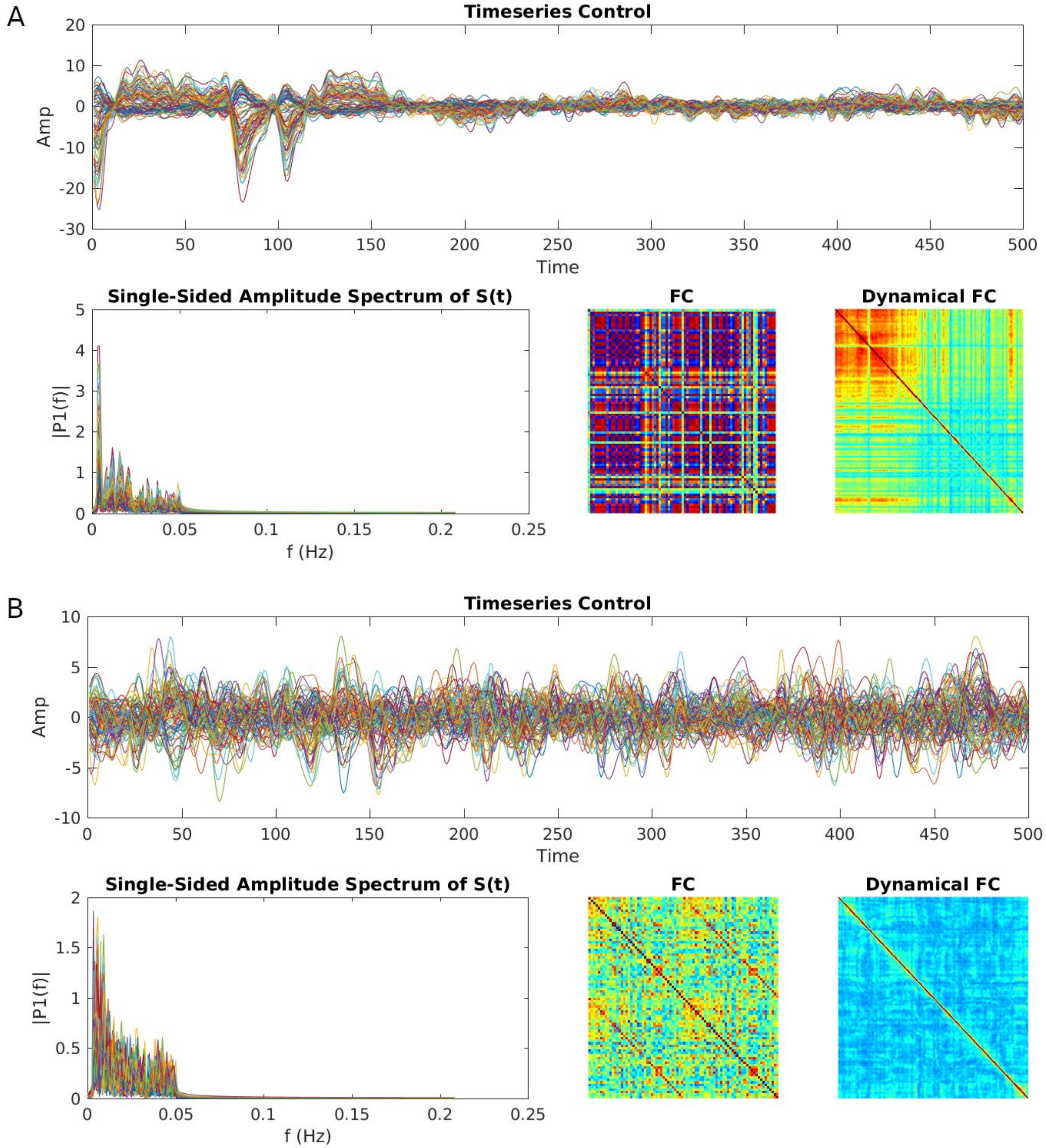

**Supplementary Figure 1. Extra Cleaning Procedure. A)** Example of discarded run for the deep sevoflurane condition. **B)** Example of a correct run for deep sevoflurane condition. In both cases, time-series, the Fourier transform, the functional connectivity and the dynamical connectivity computed with phase correlation were plotted for visual inspection. An artifact is clearly recognized in A), around 100 ms.

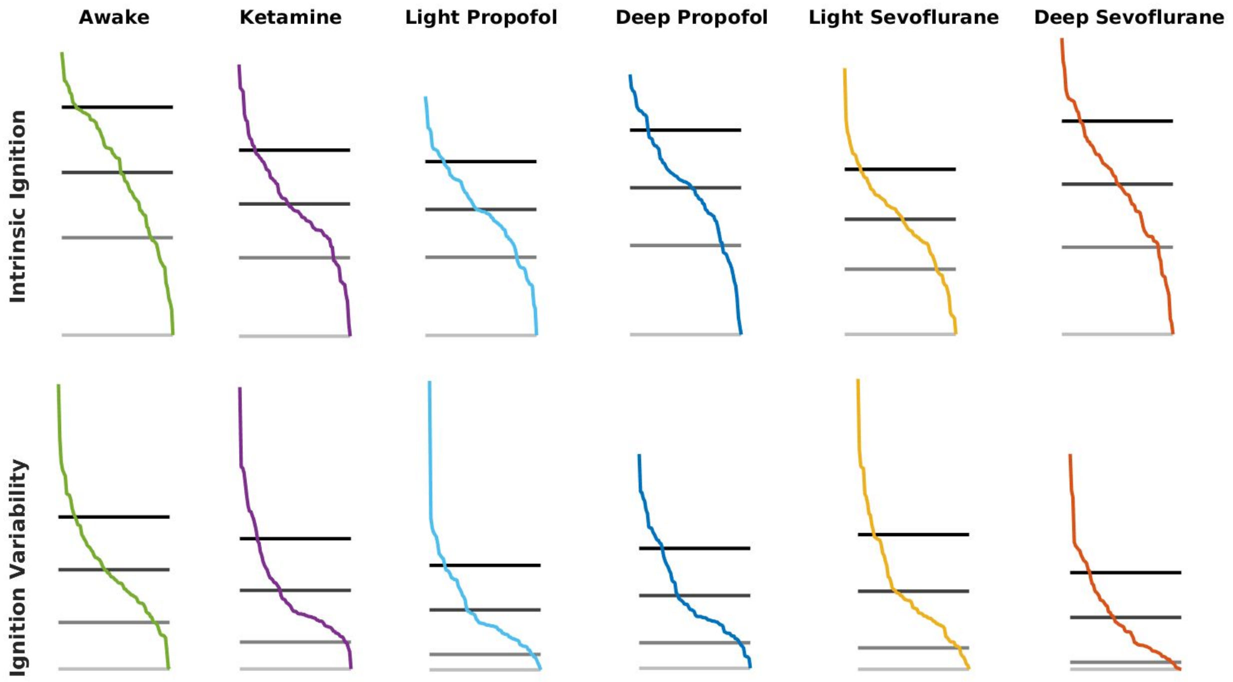

**Supplementary Figure 2. Ignition curves zoomed**. For each condition, the Intrinsic Ignition per node curve and the Ignition Variability per node curve are plotted in their own scales. Horizontal lines, from darker to lighter, correspond to $\mu+\sigma$; $\mu$; $\mu-\sigma$; $\mu-\sigma-min(node)$; $i$ is the index node, $\mu$ mean value and $\sigma$ standard deviation for each curve. The $min(node)$ is the minimum value of the curve. These curves suggest major differences in the Ignition Variability than the hierarchical structure given by the Intrinsic Ignition curve.

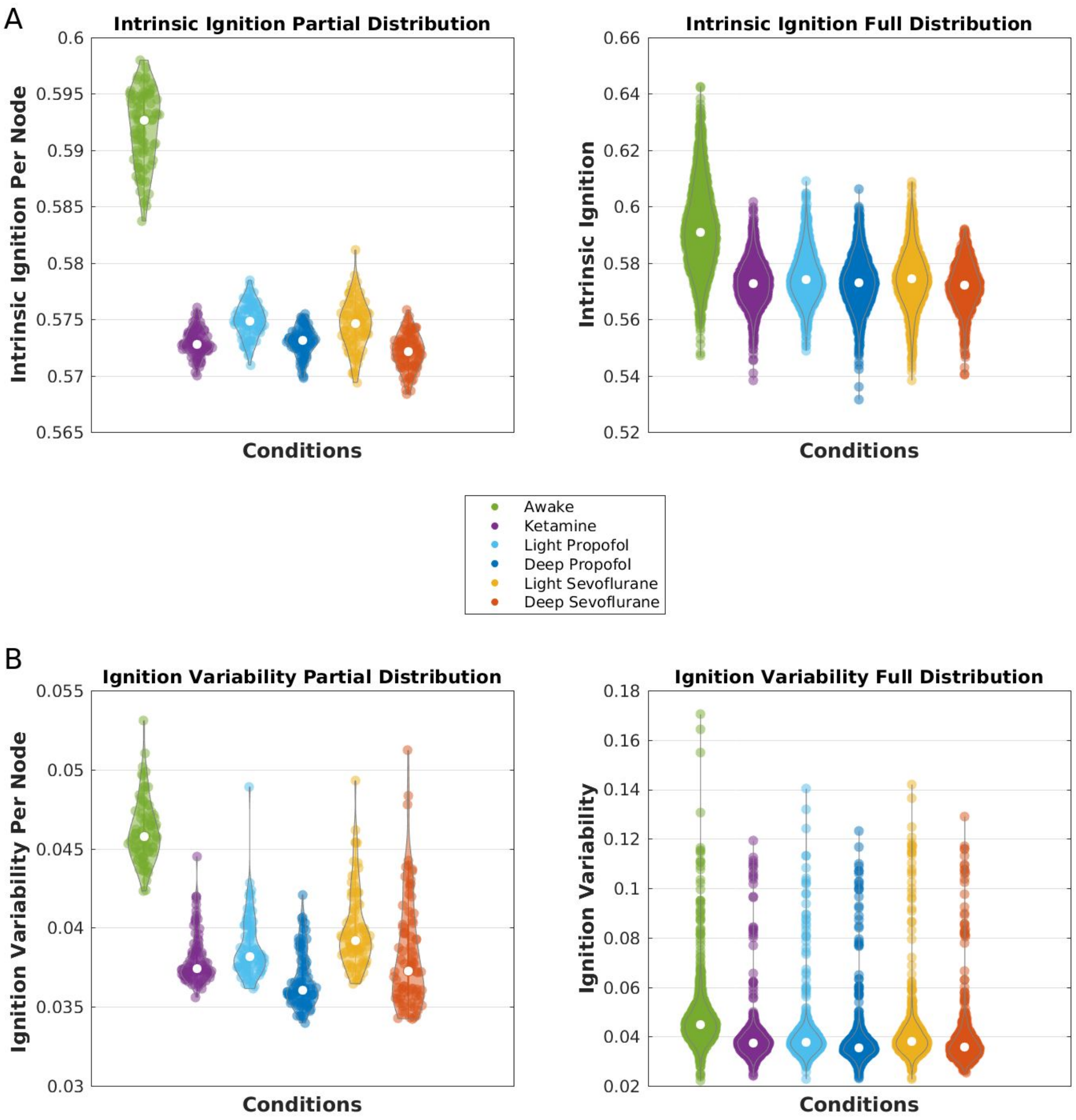

**Supplementary Figure 3. Partial versus full Density Distributions on Intrinsic Ignition.** Statistical analyses in both sets give similar results. For visualization purposes, we used the averaged data (Figure 4). **A)** Partial distribution (ROIs per condition) and Full distribution (runs x ROIs per condition) for Intrinsic Ignition. Statistically, results are similar using the partial distribution or the full distribution. Using the full distribution, awake condition differentiates from all other conditions (Kolmogorov–Smirnov test, $p<0.001$), ketamine is not significantly different than deep propofol ($p>0.01$) nor deep sevoflurane ($p>0.01$), light propofol is not different from light sevoflurane ($p>0.01$), deep propofol is not different from deep sevoflurane ($p>0.01$). The student's t-test presents a similar tendency than Kolmogorov–Smirnov test with the exception of deep propofol and dep sevoflurane (t-test, $p=0.009$). **B)** Partial distribution (ROIs per condition) and Full distribution (runs x ROIs per condition) for Ignition Variability. Using the full distribution all conditions differentiate between them (Kolmogorov–Smirnov test, $p<0.01$). Student's t-test showed a similar tendency than Kolmogorov–Smirnov test with the exception of ketamine versus deep sevoflurane (t-test, $p=0.52$), light propofol versus deep sevoflurane (t-test, $p=0.26$). Compare these results with results in Figure 4. The shape of Violin plots describes the distribution density, the white dot corresponds to the median, the thick inner line is the first quartile (down), and the third quartile (up). The borders are the upper and lower adjacent values (Hintze & Nelson, 1998).

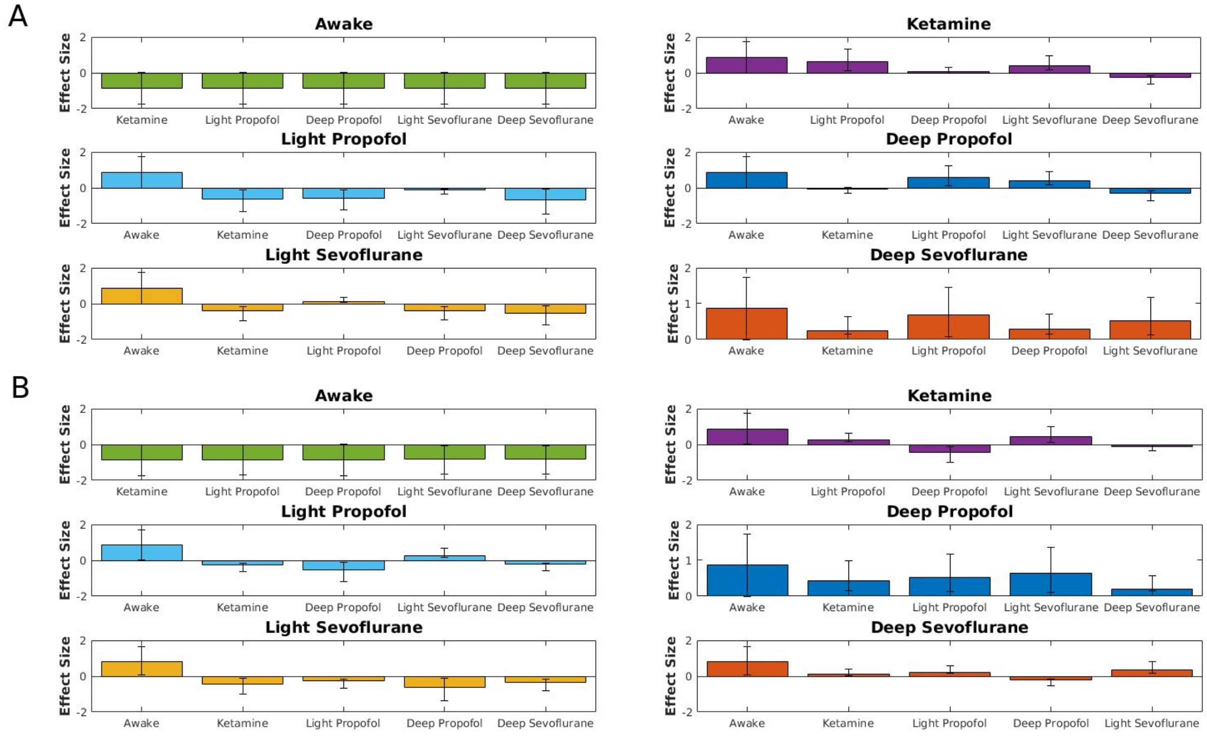

**Supplementary Figure 4. Effect Size Analysis for Intrinsic Ignition.** An effect size analysis was performed among conditions to quantify the apparent differences from the previous Figure 4 (Hentschke & Stüttgen, 2011). Bar plots show the results from the rank-biserial correlation analysis (ranks between -1 to +1 with 0 no effect) for independent samples with 10.000 bootstrapping iterations to compute confidence intervals (other effect size techniques such as mean difference, AUROC and Cohen U1 did not present major differences with the results here). **A)** The effect size analysis on the Intrinsic Ignition per node suggests bigger effects between awake and all the other sedation conditions (effect -0.866, CI [-0.8661 -0.8661]), while ketamine and deep propofol have small differences between them (effect 0.0659 CI [-0.087 0.222]), as well as light propofol in comparison with light sevoflurane (effect -0.1 CI [-0.259 0.047]. It supports previous analyses in Figure 4b. **B)** The effect size analysis on the Ignition Variability per node suggests that all the conditions have a considerable effect size, with the exception of ketamine and deep sevoflurane (effect -0.1 CI [-0.266 0.063]). Error bars are CI.

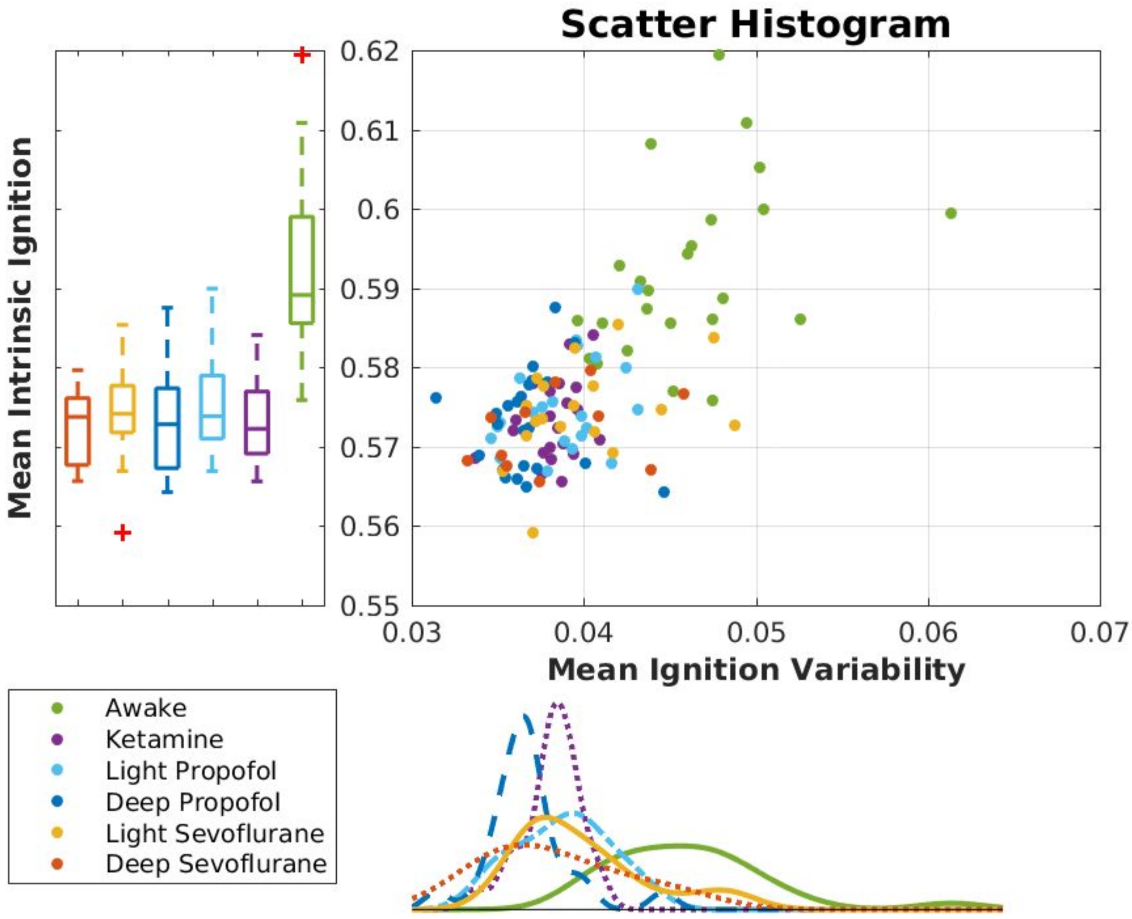

**Supplementary Figure 5. Scatter plot between Mean Intrinsic Ignition and Mean Ignition Variability.** Two clusters are not so clearly observed as before, Figure 4b. However, statistically, the mean Intrinsic Ignition in the awake condition is different from anesthetics (values in captions Figure 4d), while for the mean Ignition Variability, the awake condition is also significantly higher than sedations. Deep propofol is statistically lower than ketamine (p=0.001), light propofol (p=0.0053), and light sevoflurane (p=0.004). However other sedations are not statistically different between them (awake CI [0.0480 0.0442], ketamine CI [0.0388 0.0374], light propofol CI [0.0399 0.0377], deep propofol CI [0.0377 0.0357], light sevoflurane CI [0.0417 0.0382], deep sevoflurane CI [0.0407 0.0360])

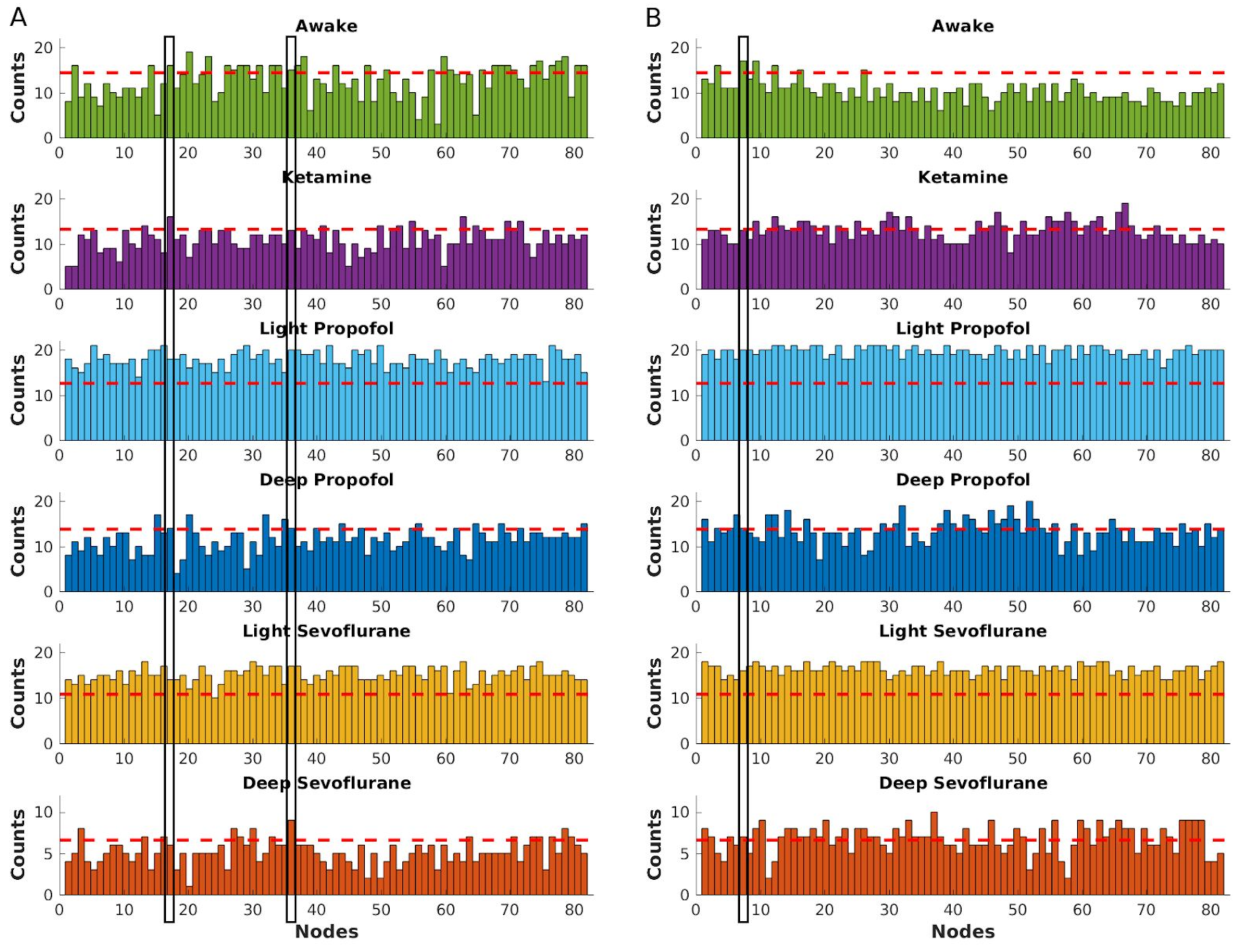

**Supplementary Figure 6. Local tendency analysis across subjects.** If the same analysis from Figures 5, 6, and Supplementary Tables 3-8 is now performed subject by subject across the six conditions, only a few nodes survived the local tendency analysis. For each subject in each condition, the logical propositions from Supplementary tables 3-8 were computed and each appearance saved to then plot the histograms for Intrinsic Ignition (**A**) and Ignition Variability (**B**). If a threshold of 60% of appearance is required (red dotted lines), only two nodes for Intrinsic Ignition and only one node for Ignition Variability satisfy the three logical propositions across the six conditions. In the first case, the regions are Subgenual cingulate cortex right and Intraparietal cortex right (index 17 and 36 respectively). In the second case, only the Central temporal cortex right (index 7) survived this restricted analysis. Black vertical squares highlight the areas mentioned.

### II. Supplementary Tables

| **Intrinsic Ignition** | **Awake** | | **Ketamine** | | **Light Propofol** | | **Deep Propofol** | | **Light Sevoflurane** | | **Deep Sevoflurane** | |
|---|---|---|---|---|---|---|---|---|---|---|---|---|
| | Effect | CI | Effect | CI | Effect | CI | Effect | CI | Effect | CI | Effect | CI |
| **Awake** | - | - | -0.87 | [-0.87 -0.87] | -0.87 | [-0.87 -0.87] | -0.87 | [-0.87 -0.87] | -0.87 | [-0.87 -0.87] | -0.87 | [-0.87 -0.87] |
| **Ketamine** | 0.87 | [0.87 0.87] | - | - | 0.61 | [0.5 0.7] | 0.66 | [-0.09 -0.22] | 0.41 | [0.27 0.54] | -0.25 | [-0.4 -0.11] |
| **Light propofol** | 0.87 | [0.87 0.87] | -0.61 | [-0.7 -0.5] | - | - | -0.57 | [-0.67 -0.46] | -0.11 | [0.26 0.04] | -0.69 | [-0.76- -0.6] |
| **Deep propofol** | 0.87 | [0.87 0.87] | -0.07 | [-0.22 0.09] | 0.57 | [0.46 0.67] | - | - | 0.4 | [0.25 0.53] | -0.29 | [-0.42 -0.14] |
| **Light sevoflurane** | 0.87 | [0.87 0.87] | -0.41 | [-0.55 -0.27] | -0.11 | [-0.05 0.26] | -0.39 | [-0.53 -0.25] | - | - | -0.53 | [-0.64- -0.4] |
| **Deep sevoflurane** | 0.87 | [0.87 0.87] | 0.25 | [0.1 0.4] | 0.69 | [0.61 0.76] | -0.29 | [0.14 0.42] | 0.53 | [0.4 0.64] | - | - |
| **Intrinsic Variability** | **Awake** | | **Ketamine** | | **Light Propofol** | | **Deep Propofol** | | **Light Sevoflurane** | | **Deep Sevoflurane** | |
| | Effect | CI | Effect | CI | Effect | CI | Effect | CI | Effect | CI | Effect | CI |
| **Awake** | - | - | -0.86 | [-0.87 -0.85] | -0.85 | [-0.87 -0.80] | -0.87 | [-0.87 -0.87] | -0.80 | [-0.84 -0.73] | -0.79 | [-0.85 -0.72] |
| **Ketamine** | 0.86 | [0.85 0.87] | - | - | 0.24 | [0.09 0.38] | 0.43 | [-0.56 -0.29] | 0.45 | [0.32 0.57] | -0.1 | [-0.27 0.06] |
| **Light propofol** | 0.85 | [0.8 0.87] | -0.24 | [-0.38 -0.08] | - | - | -0.52 | [-0.64 -0.4] | 0.26 | [0.11 0.4] | -0.2 | [-0.36 -0.04] |
| **Deep propofol** | 0.87 | [0.87 0.87] | 0.43 | [0.29 0.56] | 0.52 | [0.4 0.64] | - | - | 0.63 | [0.52 0.72] | 0.2 | [0.05 0.36] |
| **Light sevoflurane** | 0.8 | [0.73 0.84] | -0.45 | [-0.57 -0.32] | -0.26 | [-0.4 -0.11] | -0.63 | [-0.72 -0.52] | - | - | -0.33 | [-0.47 -0.18] |
| **Deep sevoflurane** | 0.79 | [0.72 0.85] | 0.1 | [-0.06 0.27] | 0.21 | [0.05 0.37] | -0.2 | [-0.35 -0.05] | 0.33 | [0.19 0.47] | - | - |

**Supplementary Table 1. Effect Size Values and Confidence intervals for Intrinsic Ignition per node and Intrinsic Variability.** Effect size analyzes quantify how much different two distributions are, instead of only focusing on if two distributions are or not different. It is a complement of common statistical tests for Figure 3a-d.

| | $i >= \mu+\sigma$ | $\mu+\sigma >= i > \mu$ | $\mu >= i > \mu-\sigma$ | $\mu-\sigma > i$ | | $i >= \mu+\sigma$ | $\mu+\sigma > i >= \mu$ | $\mu > i >= \mu-\sigma$ | $\mu-\sigma > i$ |
|---|---|---|---|---|---|---|---|---|---|
| Awake | 19<br>20<br>23<br>29<br>31<br>**38**<br>60<br>74<br>75<br>**78**<br>79 | 2<br>14<br>16<br>**17**<br>22<br>26<br>**27**<br>28<br>**33**<br>34<br>**36**<br>37<br>41<br>43<br>48<br>50<br>**51**<br>54<br>**58**<br>61<br>62<br>63<br>64<br>66<br>68<br>69<br>70<br>**71**<br>72<br>**77**<br>81<br>82 | 3<br>4<br>7<br>10<br>11<br>13<br>21<br>25<br>30<br>32<br>35<br>40<br>42<br>44<br>45<br>46<br>49<br>52<br>55<br>57<br>67<br>73 | 1<br>5<br>6<br>8<br>9<br>12<br>15<br>18<br>24<br>39<br>47<br>53<br>56<br>59<br>65<br>80 | Ketamine | 1<br>2<br>8<br>9<br>20<br>28<br>45<br>47<br>48<br>56<br>69<br>73 | 6<br>18<br>19<br>21<br>24<br>29<br>32<br>35<br>39<br>42<br>44<br>46<br>49<br>54<br>59<br>60<br>61<br>62<br>64<br>66<br>67<br>68<br>75<br>79 | 3<br>4<br>7<br>10<br>11<br>12<br>13<br>14<br>15<br>16<br>23<br>25<br>**27**<br>**33**<br>**36**<br>37<br>**38**<br>40<br>41<br>50<br>**51**<br>52<br>53<br>55<br>**58**<br>65<br>**71**<br>74<br>76<br>**77**<br>**78**<br>80<br>82 | 5<br>**17**<br>22<br>26<br>30<br>31<br>34<br>43<br>57<br>63<br>70<br>72<br>81 |
| Light Propofol | 2<br>3<br>8<br>12<br>24<br>26<br>35<br>41<br>43<br>45<br>65<br>76<br>79<br>82 | 4<br>5<br>10<br>11<br>14<br>18<br>20<br>23<br>28<br>31<br>**38**<br>39<br>**51**<br>52<br>53<br>54<br>56<br>**58**<br>60<br>61<br>64 | 1<br>6<br>7<br>9<br>13<br>16<br>**17**<br>19<br>21<br>22<br>25<br>29<br>30<br>**33**<br>34<br>**36**<br>44<br>47<br>49<br>57<br>59 | 15<br>**27**<br>32<br>37<br>40<br>42<br>46<br>48<br>50<br>55<br>68<br>70<br>73<br>75<br>**77** | Deep Propofol | 1<br>6<br>13<br>14<br>18<br>19<br>23<br>25<br>29<br>48<br>52<br>59<br>64<br>67 | 2<br>3<br>4<br>5<br>7<br>8<br>11<br>16<br>22<br>26<br>31<br>34<br>39<br>40<br>41<br>43<br>45<br>49<br>50<br>53<br>55 | 9<br>12<br>21<br>24<br>**27**<br>28<br>30<br>**33**<br>35<br>**36**<br>37<br>**38**<br>44<br>46<br>54<br>**58**<br>62<br>65<br>70<br>**71**<br>73 | 10<br>15<br>**17**<br>20<br>32<br>42<br>47<br>**51**<br>56<br>57<br>66<br>69<br>72<br>**78** |

| | | | | | | | | | |
|---|---|---|---|---|---|---|---|---|---|
| | | 66<br>72<br>80 | 62<br>63<br>67<br>69<br>**71**<br>74<br>**78**<br>81 | | | | 60<br>61<br>63<br>68<br>74<br>76<br>79<br>80<br>81<br>82 | 75<br>**77** | |
| Light Sevoflurane | 2<br>4<br>5<br>20<br>24<br>28<br>35<br>39<br>47<br>48<br>49<br>61 | 7<br>8<br>10<br>11<br>12<br>14<br>15<br>**17**<br>18<br>19<br>21<br>26<br>**38**<br>41<br>43<br>46<br>52<br>53<br>56<br>57<br>**58**<br>59<br>60<br>64<br>65<br>67<br>68<br>69<br>76<br>80<br>82 | 1<br>6<br>9<br>16<br>22<br>23<br>25<br>**27**<br>34<br>40<br>44<br>45<br>50<br>**51**<br>55<br>62<br>66<br>70<br>**71**<br>73<br>**77**<br>**78**<br>79<br>81 | 3<br>13<br>29<br>30<br>31<br>32<br>**33**<br>**36**<br>37<br>42<br>54<br>63<br>72<br>74<br>75 | Deep Sevoflurane | 1<br>5<br>6<br>14<br>20<br>23<br>25<br>31<br>46<br>48<br>50<br>53<br>57<br>70 | 4<br>7<br>11<br>15<br>18<br>19<br>22<br>26<br>32<br>35<br>37<br>39<br>40<br>41<br>42<br>45<br>54<br>56<br>59<br>60<br>61<br>62<br>63<br>65<br>66<br>67<br>72<br>76 | 2<br>3<br>9<br>10<br>12<br>13<br>16<br>**17**<br>21<br>24<br>30<br>**38**<br>43<br>44<br>47<br>49<br>**51**<br>52<br>55<br>64<br>68<br>**71**<br>73<br>74<br>75<br>**78**<br>79<br>81<br>82 | 8<br>**27**<br>28<br>29<br>**33**<br>34<br>**36**<br>**58**<br>69<br>**77**<br>80 |

**Supplementary Table 2. Names and indexes region of Interest in CoCoMac parcellation Intrinsic Ignition all Condition.** Red bold indexes correspond to the nodes that satisfy the three conditions: $nodes \geq \mu$ for awake, $\mu + \sigma \geq nodes$ for light conditions and $\mu \geq nodes$ for deep conditions.

| Awake | $i >= \mu + \sigma$ | $\mu + \sigma >= i > \mu$ | $\mu >= i > \mu - \sigma$ | $\mu - \sigma > i$ | Ketamine | $i >= \mu + \sigma$ | $\mu + \sigma > i >= \mu$ | $\mu > i >= \mu - \sigma$ | $\mu - \sigma > i$ |
|---|---|---|---|---|---|---|---|---|---|
| | 3<br>6<br>**7**<br>9<br>12 | **1**<br>10<br>16<br>21<br>26 | 2<br>4<br>5<br>8<br>11 | 19<br>23<br>31<br>33<br>34 | | 6<br>15<br>23<br>26<br>27 | 8<br>11<br>16<br>21<br>24 | **1**<br>2<br>3<br>4<br>5 | 9<br>29<br>55<br>63<br>67 |

| | | | | | | | | | |
|---|---|---|---|---|---|---|---|---|---|
| | 13<br>14<br>**17**<br>37<br>43<br>48<br>53<br>78 | 27<br>30<br>35<br>40<br>47<br>50<br>51<br>54<br>55<br>59<br>62<br>72<br>74<br>**76**<br>79<br>82 | 15<br>18<br>20<br>22<br>24<br>25<br>28<br>29<br>32<br>36<br>38<br>39<br>41<br>42<br>44<br>49<br>52<br>56<br>58<br>60<br>61<br>64<br>65<br>66<br>67<br>68<br>69<br>71<br>73<br>80<br>81 | 45<br>46<br>57<br>63<br>70<br>75<br>77 | | 41<br>42<br>43<br>53<br>70<br>80<br>81<br>82 | 30<br>31<br>34<br>35<br>40<br>48<br>49<br>57<br>61<br>65<br>69<br>75 | **7**<br>10<br>12<br>13<br>14<br>**17**<br>18<br>19<br>20<br>22<br>25<br>28<br>32<br>33<br>36<br>37<br>38<br>39<br>44<br>45<br>46<br>47<br>50<br>51<br>52<br>54<br>56<br>58<br>59<br>60<br>62<br>64<br>66<br>68<br>71<br>72<br>73<br>74<br>**76**<br>77<br>78<br>79 | |
| Light Propofol | 32<br>40<br>48<br>49<br>51<br>53<br>55<br>57<br>59<br>61<br>72<br>73 | 5<br>9<br>10<br>11<br>14<br>23<br>37<br>38<br>46<br>56<br>67<br>68<br>69<br>71<br>74<br>78 | **1**<br>2<br>3<br>4<br>6<br>**7**<br>8<br>12<br>13<br>15<br>16<br>18<br>19<br>20<br>21<br>24<br>25<br>27<br>28<br>29<br>30<br>31 | **17**<br>22<br>26<br>58<br>62<br>63 | Deep Propofol | 3<br>9<br>10<br>15<br>19<br>20<br>23<br>36<br>54<br>56<br>57<br>60<br>64<br>69<br>72<br>75<br>78 | 8<br>13<br>14<br>16<br>26<br>33<br>37<br>42<br>47<br>51<br>62<br>77 | **1**<br>2<br>4<br>5<br>**7**<br>12<br>**17**<br>18<br>21<br>22<br>24<br>25<br>27<br>28<br>29<br>30<br>31<br>34<br>35<br>38<br>39<br>40 | 6<br>11<br>32<br>43<br>49<br>52<br>53<br>59 |

| | | | | | | | | | |
|---|---|---|---|---|---|---|---|---|---|
| | | | 33<br>34<br>35<br>36<br>39<br>41<br>42<br>43<br>44<br>45<br>47<br>50<br>52<br>54<br>60<br>64<br>65<br>66<br>70<br>75<br>**76**<br>77<br>79<br>80<br>81<br>82 | | | | | 41<br>44<br>45<br>46<br>48<br>50<br>55<br>58<br>61<br>63<br>65<br>66<br>67<br>68<br>70<br>71<br>73<br>74<br>**76**<br>79<br>80<br>81<br>82 | |
| Light Sevoflurane | 23<br>29<br>30<br>31<br>40<br>50<br>53<br>69<br>70<br>73<br>74<br>79 | 6<br>20<br>25<br>32<br>33<br>36<br>51<br>55<br>59<br>65<br>66<br>67<br>71<br>75<br>80<br>81 | 2<br>3<br>4<br>5<br>**7**<br>8<br>10<br>11<br>12<br>13<br>14<br>15<br>16<br>**17**<br>18<br>19<br>21<br>22<br>24<br>26<br>27<br>28<br>34<br>35<br>37<br>38<br>39<br>41<br>42<br>43<br>44<br>45<br>46<br>47<br>48<br>49<br>52<br>56 | **1**<br>9<br>54<br>61<br>64<br>**76**<br>77 | Deep Sevoflurane | 2<br>8<br>11<br>12<br>24<br>27<br>34<br>43<br>49<br>52<br>53<br>58<br>69<br>80<br>81 | 4<br>5<br>21<br>28<br>29<br>30<br>33<br>40<br>41<br>44<br>51<br>55<br>61<br>68<br>71<br>72<br>75 | **1**<br>3<br>6<br>**7**<br>9<br>13<br>14<br>15<br>16<br>**17**<br>18<br>19<br>22<br>23<br>25<br>31<br>32<br>36<br>38<br>39<br>42<br>45<br>47<br>48<br>50<br>54<br>56<br>57<br>59<br>60<br>62<br>63<br>64<br>65<br>66<br>67<br>70<br>73 | 10<br>20<br>26<br>35<br>37<br>46<br>77<br>78 |

| | | | | | | | | |
|---|---|---|---|---|---|---|---|---|
| | | | 57<br>58<br>60<br>62<br>63<br>68<br>72<br>78<br>82 | | | | | 74<br>**76**<br>79<br>82 |

**Supplementary Table 3. Names and indexes region of Interest in CoCoMac parcellation Ignition Variability in all Condition.** Red bold indexes correspond to the nodes that satisfy the three conditions: $nodes \geq \mu$ for awake, $\mu + \sigma \geq nodes$ for light conditions and $\mu \geq nodes$ for deep conditions.

| Index | Name | Hemisfere | Acronyms |
|---|---|---|---|
| 1 | Tempolar polar | Right | TCpol |
| 2 | Superior temporal cortex | Right | TCs |
| 3 | Amygdala | Right | Amyg |
| 4 | Orbitoinferior prefrontal cortex | Right | PFCoi |
| 5 | Anterior insula | Right | Ia |
| 6 | Orbitomedial prefrontal cortex | Right | PFCom |
| 7 | Central temporal cortex | Right | TCc |
| 8 | Orbitolateral prefrontal cortex | Right | PFCol |
| 9 | Inferior temporal | Right | TCi |
| 10 | Parahippocampal cortex | Right | PHC |
| 11 | Gustatory cortex | Right | G |
| 12 | Ventrolateral premotor cortex | Right | PMCvl |
| 13 | Anterior visual area (ventral) | Right | VACv |
| 14 | Posterior insula | Right | Ip |
| 15 | Prefrontal polar cortex | Right | PFCpol |
| 16 | Hippocampus | Right | HC |
| 17 | Subgenual cingulate cortex | Right | CCs |
| 18 | Ventrolateral prefrontal cortex | Right | PFCvl |
| 19 | Visual area 2 | Right | V2 |
| 20 | Medial prefrontal cortex | Right | PFCm |

| | | | |
|---|---|---|---|
| 21 | Ventral temporal cortex | Right | TCv |
| 22 | Anterior visual area (dorsal) | Right | VACd |
| 23 | Visual area 1 | Right | V1 |
| 24 | Centrolateral prefrontal cortex | Right | PFCcl |
| 25 | Secondary auditory cortex | Right | A2 |
| 26 | Retrosplenial cingulate cortex | Right | CCr |
| 27 | Posterior cingulate cortex | Right | CCp |
| 28 | Anterior cingulate cortex | Right | CCa |
| 29 | Secondary somatosensory cortex | Right | S2 |
| 30 | Primary somatosensory cortex | Right | S1 |
| 31 | Primary auditory cortex | Right | A1 |
| 32 | Primary motor cortex | Right | M1 |
| 33 | Inferior parietal cortex | Right | PCi |
| 34 | Medial parietal cortex | Right | PCm |
| 35 | Dorsomedial prefrontal cortex | Right | PFCdm |
| 36 | Intraparietal cortex | Right | PCip |
| 37 | Superior parietal cortex | Right | PCs |
| 38 | Frontal eye field | Right | FEF |
| 39 | Dorsolateral prefrontal cortex | Right | PFCdl |
| 40 | Medial premotor cortex | Right | PMCm |
| 41 | Dorsolateral premotor cortex | Right | PMCdl |
| 42 | Tempolar polar | Left | TCpol |
| 43 | Superior temporal cortex | Left | TCs |
| 44 | Amygdala | Left | Amyg |
| 45 | Orbitoinferior prefrontal cortex | Left | PFCoi |
| 46 | Anterior insula | Left | Ia |
| 47 | Orbitomedial prefrontal cortex | Left | PFCom |
| 48 | Central temporal cortex | Left | TCc |
| 49 | Orbitolateral prefrontal cortex | Left | PFCol |
| 50 | Inferior temporal | Left | TCi |

| | | | |
|---|---|---|---|
| 51 | Parahippocampal cortex | Left | PHC |
| 52 | Gustatory cortex | Left | G |
| 53 | Ventrolateral premotor cortex | Left | PMCvl |
| 54 | Anterior visual area (ventral) | Left | VACv |
| 55 | Posterior insula | Left | Ip |
| 56 | Prefrontal polar cortex | Left | PFCpol |
| 57 | Hippocampus | Left | HC |
| 58 | Subgenual cingulate cortex | Left | CCs |
| 59 | Ventrolateral prefrontal cortex | Left | PFCvl |
| 60 | Visual area 2 | Left | V2 |
| 61 | Medial prefrontal cortex | Left | PFCm |
| 62 | Ventral temporal cortex | Left | TCv |
| 63 | Anterior visual area (dorsal) | Left | VACd |
| 64 | Visual area 1 | Left | V1 |
| 65 | Centrolateral prefrontal cortex | Left | PFCcl |
| 66 | Secondary auditory cortex | Left | A2 |
| 67 | Retrosplenial cingulate cortex | Left | CCr |
| 68 | Posterior cingulate cortex | Left | CCp |
| 69 | Anterior cingulate cortex | Left | CCa |
| 70 | Secondary somatosensory cortex | Left | S2 |
| 71 | Primary somatosensory cortex | Left | S1 |
| 72 | Primary auditory cortex | Left | A1 |
| 73 | Primary motor cortex | Left | M1 |
| 74 | Inferior parietal cortex | Left | PCi |
| 75 | Medial parietal cortex | Left | PCm |
| 76 | Dorsomedial prefrontal cortex | Left | PFCdm |
| 77 | Intraparietal cortex | Left | PCip |
| 78 | Superior parietal cortex | Left | PCs |
| 79 | Frontal eye field | Left | FEF |
| 80 | Dorsolateral prefrontal cortex | Left | PFCdl |

| | | | |
|---|---|---|---|
| 81 | Medial premotor cortex | Left | PMCm |
| 82 | Dorsolateral premotor cortex | Left | PMCdl |

**Supplementary Table 4. Names and indexes region of Interest in CoCoMac parcellation.**
